# 40 Hz sensory stimulation enhances CA3-CA1 coordination and prospective coding during navigation in a mouse model of Alzheimer’s disease

**DOI:** 10.1101/2024.10.23.619408

**Authors:** Abigail L. Paulson, Lu Zhang, Ashley M. Prichard, Annabelle C. Singer

**Affiliations:** Coulter Department of Biomedical Engineering, Emory University and Georgia Institute of Technology, Atlanta, GA, 30332, USA; National Institute of Mental Health, NIH, Bethesda, 20892, MD; Atlanta VA Medical Center, Decatur, GA, 30033, USA

**Keywords:** Alzheimer’s disease, Spatial navigation, Hippocampus, Non-invasive brain stimulation, Prospective Coding, Replay, Gamma oscillations

## Abstract

40 Hz sensory stimulation (“flicker”) has emerged as a new technique to potentially mitigate pathology and improve cognition in mouse models of Alzheimer’s disease (AD) pathology. However, it remains unknown how 40 Hz flicker affects neural codes essential for memory. Accordingly, we investigate the effects of 40 Hz flicker on neural representations of experience in the hippocampus of the 5XFAD mouse model of AD by recording 1000s of neurons during a goal-directed spatial navigation task. We find that an hour of daily exposure to 40 Hz audio-visual stimulation over 8 days leads to higher coordination between hippocampal subregions CA3 and CA1 during navigation. Consistent with CA3’s role in generating sequential activity that represents future positions, 40 Hz flicker exposure increased prospective coding of future positions. In turn, prospective coding was more prominent during efficient navigation behavior. Our findings show how 40 Hz flicker enhances key hippocampal activity during behavior that is important for memory.

**Significance Statement:** Brain stimulation has emerged as a new potential therapeutic approach to potentially correct or improve altered neural activity in Alzheimer’s disease. One such approach, 40 Hz sensory stimulation, or flicker, has been shown to improve cognition in disease models. However, it is not clear how 40 Hz flicker affects neural activity underlying memory processes. Here, we investigate how 40 Hz flicker exposure affects neural activity patterns that are crucial for memory. We find 40Hz flicker increases neural coordination in memory circuits, indicating better communication. Furthermore, 40Hz flicker increased neural representations of future positions, patterns theorized to support memory-based planning. These results indicate that 40 Hz flicker increases key neural activity that is important for memory.

## Introduction

Manipulating neural activity has emerged as an exciting potential therapeutic approach for Alzheimer’s disease (AD), the most common dementia. Aberrant neural activity occurs early in AD progression and is thought to play a key role in both molecular pathology and cognitive impairment (Bero et al., 2011; Koenig et al., 2005; Palmqvist et al., 2017; Sorg et al., 2007). Gamma oscillations, 30-100 Hz electrical oscillations that coordinate neural activity, have been of special interest because they support cognition by coordinating neural activity and are altered in humans with AD, mouse models of AD pathology, and models of genetic risk of AD (Fries et al., 2007; Gillespie et al., 2016; Iaccarino et al., 2016; Stam et al., 2002; Verret et al., 2012). Rescuing aspects of neural activity involved in the generation of gamma reduces AD-associated pathology and improves cognition in mouse models (Gillespie et al., 2016; Iaccarino et al., 2016; Martinez-Losa et al., 2018; Verret et al., 2012). As a result, targeting gamma activity as a potential intervention in AD is actively under investigation. Recent work shows stimulating 40 Hz activity, which is within the gamma range, via optogenetic stimulation reduces amyloid beta, a peptide thought to initiate a neurotoxic cascade in AD, and recruits microglia to engulf amyloid (Iaccarino et al., 2016). These effects on amyloid beta and microglia have also been achieved by exposing animals to 40 Hz flicker, lights and sounds flashing at 40 Hz, that have been referred to as gamma sensory stimulation, 40 Hz audio or audio-visual stimulation, GENUS, or multisensory flicker (Iaccarino et al., 2016; Martorell et al., 2019; Park et al., 2020, 2022; Shen et al., 2022). Exposing a mouse model of amyloidosis to 7 days of 40 Hz audio or audio-visual flicker improves performance on spatial and working memory tasks (Martorell, et al., 2019). These results have inspired further work revealing the intracellular signaling by which 40 Hz flicker affects amyloid beta processing, as well as interactions between flicker and exercise in AD, the effects of flicker in other disease models and human patients, and the effects of other forms of gamma frequency stimulation (Adaikkan et al., 2019; Benussi et al., 2022; Blanpain et al., 2024; He et al., 2021; Kim et al., 2022; Liu et al., 2023; Murdock et al., 2024; Park et al., 2020, 2022; Shen et al., 2022; Sprugnoli et al., 2021; Yao et al., 2020; Zheng et al., 2020). However, the mechanisms through which 40 Hz flicker acts on memory processes are still unknown. While endogenous gamma has long been associated with coordinating neural activity within and across regions to enhance communication and neural codes, 40 Hz flicker is different from endogenous gamma and it is unclear how flicker affects such neural coordination (Fries, 2009). Understanding how flicker acts on neural communication and neural codes is essential to developing flicker as a research and clinical tool. Thus, because 40 Hz flicker exposure improves cognitive function in mouse models of AD, we investigated how 40 Hz flicker affects neural activity that is known to be important for learning and memory.

In the hippocampus, a region crucial for spatial and episodic learning and memory that is affected early in AD, neural activity represents ongoing and previous experiences to support memory encoding and retrieval (Braak & Braak, 1991; Eichenbaum, 2004). Gamma activity is thought to play a role in these cognitive processes by coordinating neurons to fire together on short timescales, supporting communication and information transfer between brain regions (Fries et al., 2007). When two regions oscillate at the same frequency with the right relative timing, inputs from one region arrive when the downstream target neurons are most depolarized and likely to fire in response, thus creating strong functional connectivity, called communication via coherence (Fries, 2005). Prior work has shown that coupling between CA3 and CA1 subfields of the hippocampus at slow gamma (often defined as 30-50 Hz or 25-55 Hz) oscillatory frequencies enhances intrahippocampal communication to facilitate encoding and memory retrieval (Colgin et al., 2009; Montgomery & Buzsáki, 2007). Indeed, increased slow gamma coherence between CA3 and CA1 during encoding was shown to predict subsequent successful spatial memory performance (Trimper et al., 2014, 2017). Slow gamma is altered in multiple models of AD and reduced slow gamma in CA3 has been shown to predict future memory impairment, suggesting that slow gamma disruptions in the hippocampal circuit contribute to cognitive deficits (Gillespie et al., 2016; Iaccarino et al., 2016; Jones et al., 2019; Verret et al., 2012).

We hypothesized that 40 Hz flicker affects neural representations that are important for memory, especially representations of past and future experiences. CA3 generates sequences of activity that represent past, current, and future experiences, and the generation of these sequences and their propagation to CA1, the primary output region of hippocampus, is crucial for memory processes (Middleton & McHugh, 2016; Nakashiba et al., 2009). During theta oscillations (4-10 Hz activity observed during running), a subset of hippocampal pyramidal cells, termed place cells, fire when the animal is in a specific location in space. While traversing an environment during navigation behavior, sequences of place cells fire, representing spatial trajectories or paths through the environment (Foster & Wilson, 2007; O’Keefe, 1976; Skaggs et al., 1996). These patterns, called theta sequences, carry information about the animal’s current position along with near future and past locations on the compressed timescales required for plasticity. CA3 inputs and associated slow gamma are essential for coordinating these sequences of activity that represent future locations, or prospective coding (Bieri et al., 2014; Middleton & McHugh, 2016). Prospective coding also occurs in bursts of hippocampal population activity during sharp-wave ripples (SWRs), 150-250 Hz oscillations essential for learning and memory that occur during pauses in behavior and sleep (Fernández-Ruiz et al., 2019; Girardeau et al., 2009; Jadhav et al., 2012). During SWRs, long trajectories of future and past locations are reactivated. Increased slow gamma power during SWRs is correlated with higher fidelity reactivation of these sequences, meaning the neurons’ firing order during the SWR better reflects sequences of positions in the environment (Carr et al., 2012; Cheng & Frank, 2008; Foster & Wilson, 2006; Wilson & McNaughton, 1994). SWRs, slow gamma during SWRs, and associated spiking activity are deficient in models of AD and may contribute to cognitive impairment since they are essential for spatial learning and memory (Fernández-Ruiz et al., 2019; Gillespie et al., 2016; Girardeau et al., 2009; Iaccarino et al., 2016; Jadhav et al., 2012; Jones et al., 2019; Prince et al., 2021). Despite the role of these neural codes in memory and their known deficits in AD, it is unknown how neural stimulation interventions like 40 Hz flicker affect this important hippocampal activity.

The neural mechanisms underlying 40 Hz flicker have been debated, with key questions raised about how 40 Hz flicker affects hippocampal activity, particularly endogenous hippocampal gamma (Duecker et al., 2021; Soula et al., 2023). 40 Hz flicker was inspired by deficits in gamma, but it is unclear if flicker has similar effects in the brain as endogenous slow gamma activity (Soula et al., 2023). Thus, further study of flicker’s effects on hippocampal neural activity is sorely needed. Prior work shows that CA3 is a strong source of hippocampal slow gamma suggesting that connections between CA3 interneurons and pyramidal cells generate slow gamma in the hippocampus (Csicsvari et al., 2003). As a result, the intrinsic properties of this circuit may make CA3 especially well-suited to respond to a 40 Hz stimulus. Indeed, one study shows that 40 Hz flicker induces synaptic plasticity between CA3 and CA1 that may lead to long term changes in the CA3-CA1 circuit (Zheng et al., 2020). We hypothesized that stimulating 40 Hz activity would elicit key similar effects in hippocampus as endogenous slow gamma, promoting stronger CA3-CA1 communication and prospective codes, even without directly entraining endogenous oscillations. Specifically, we hypothesized that 40 Hz flicker increases coordinated neural activity between CA3 and CA1, resulting in increased prospective codes (**Figure 1A**). However, given that any stimulation is artificial, 40 Hz flicker may have very different effects than endogenous rhythms within the brain. Recent studies suggest that responses to 40 Hz flicker stimulation co-exist with endogenous rhythms, highlighting the need to further investigate any long-lasting circuit effects of flicker stimulation (Duecker et al., 2021). Understanding how 40 Hz flicker stimulation affects hippocampal activity that underlies memory processes is crucial to evaluating and optimizing this technique as a potential therapeutic.

**Figure 1.**
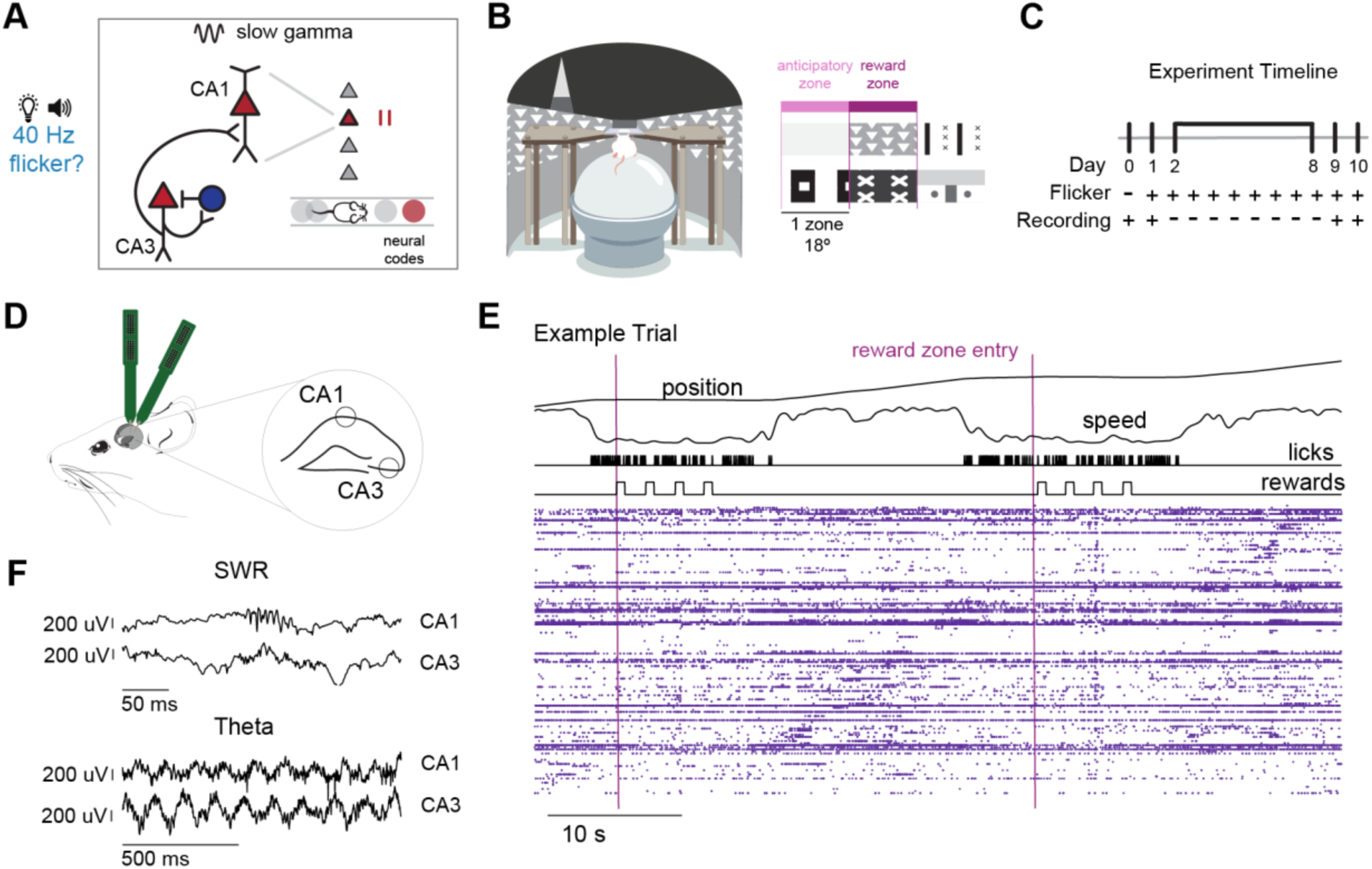
5XFAD mice perform a goal-directed spatial navigation task before and after 8 days of 40 Hz flicker. A. Hypothesized effects of 40 Hz sensory stimulation, or “flicker,” in the hippocampal circuit. Slow gamma (top, black lines) promotes CA3-CA1 communication (left) and coordinates firing patterns that code for future positions, or prospective coding (right), both of which are important for memory. We tested the hypothesis that chronic 40 Hz flicker (right) has similar effects on hippocampal function after stimulation, resulting in increased CA3-CA1 communication and prospective coding. Red triangles represent excitatory pyramidal cells, and blue circles represent inhibitory interneurons in the circuit schematic. Grey and red circles show pyramidal cells’ place fields as the mouse navigates with the red place field representing a future position. B. 6–7-month-old male 5XFAD mice were trained to run head-fixed on a spherical treadmill (left) through a continuous annular virtual-reality environment. Distinct visual cues tiled the track, and animals learned to lick in two specific patterned zones to receive a sweetened condensed milk reward (right, cues for anticipatory zones, reward zones, and one zone after are shown). (See Additional File 1 for a movie of the task.) C. Following behavior training (see Methods), electrophysiology data collection commenced. The experiment included two recording days, 7 days of flicker exposure without recordings, followed by two recording days. During recording on Experiment Day 0, animals performed the VR task only and were not exposed to flicker. During recording on Experiment Days 1, 9, and 10, animals performed the VR task and were exposed to flicker in separate sessions. D. Neural activity was recorded from dorsal hippocampal areas CA1 and CA3 simultaneously, using two multi-shanked high-density silicon electrodes. E. Neural activity was recorded as animals performed the VR spatial navigation task. *Top*, behavioral metrics recorded, including position (degrees), speed (normalized units), lick times, and reward delivery. Pink lines indicate entry into the reward zones. After a lick in the reward zone was detected, 4 rewards were delivered which encouraged pausing in the reward zone to assess SWR activity. *Bottom*, spike times (purple dots) from single neurons in CA3 and CA1 during the behavioral trial above. Each row is a separate cell including both putative pyramidal cells and interneurons. Cells are sorted by peak firing location in the track. F. Example sharp-wave ripple event (top) and example theta traces (bottom) recorded from hippocampal areas CA1 and CA3.

Here, we determined the effects of 40 Hz flicker on hippocampal neural activity that is essential for learning and memory. We investigated how chronic exposure to either 40 Hz or randomized (Random, sham condition) flicker affected hippocampal activity in 5XFAD mice during a goal-directed virtual reality spatial navigation task. We found that, compared to Random flicker, one hour of 40 Hz flicker per day over 8 days increased coupling between hippocampal subregions CA3 and CA1 during navigation behavior. Given the role of CA3-CA1 coupling on coding in hippocampus, we assessed neural codes during navigation behavior and found increased prospective coding during theta sequences and SWRs, while properties of CA3 and CA1 place cells and SWR duration and abundance remained unchanged. Because prospective codes represent future information and are thought to be important for planning ahead during navigation, and because prior work shows that the representation of forward locations increases with higher running speeds, we examined how prospective codes relate to efficient task performance (Gupta et al., 2012; Maurer et al., 2012; Parra-Barrero et al., 2021). We found prospective coding during theta sequences was increased during more efficient trials with higher running speeds, specifically in the areas of the track immediately following the reward zones when prospective coding may be advantageous in executing the next trial. Furthermore, after 40 Hz flicker, prospective coding is higher when animals lick more in anticipation of reward and on correct trials when animals were engaged in the task as compared to incorrect unengaged trials. Importantly, 40 Hz flicker induced increases in prospective coding relative to Random stimulation that were not explained by differences in speed. These findings show that exposure to 40 Hz flicker increases CA3-CA1 hippocampal coordination and prospective neural codes, suggesting possible mechanisms through which 40 Hz flicker may act in the hippocampal circuit to improve memory.

## Results

### 40 Hz flicker increases CA3-CA1 functional coupling

To test the effects of 40 Hz sensory stimulation on hippocampal neural activity important for learning and memory, we trained animals to perform a goal-directed virtual reality (VR) spatial navigation task. In this task, 6-7-month-old male 5XFAD mice learned to run through a continuous annular VR environment while head fixed on a spherical treadmill. The genotype, age, and sex of the animals was chosen to match prior work that showed gamma sensory stimulation improves spatial memory (Martorell et al., 2019). Mice learned to lick to receive a sweetened condensed reward in two areas of the track, indicated by distinct visual cues (**Figure 1B, Additional File 1**). Animals in both groups displayed anticipatory behavior, beginning to lick and reduce their speed when approaching the reward zone (“anticipatory zone”, zone immediately before the reward zone) (**Figure 1B, Supplementary Figure 1C, F, H**). After behavior training, hippocampal neural activity was recorded prior to and immediately following 8 days of audio-visual sensory stimulation (“prolonged exposure”, **Figure 1C**). On experiment day 0, animals performed the VR task only, flicker was administered on days 1 through 10, and on experiment days 1, 8, and 9 animals performed the VR task before and following 1 hour of audio-visual flicker. Animals were exposed to audio-visual flicker delivered periodically at 40 Hz (40 Hz flicker) or with randomized inter-pulse intervals with a mean of 40 Hz (Random flicker, control, see Methods). 40 Hz frequency stimulation was used because it was previously found to improve spatial memory performance compared to Random or no flicker(Martorell et al., 2019). Random flicker controlled for key aspects of the stimulus, so that the two groups were exposed to a similar amount of light and sound pulses during the stimulation, but only the 40 Hz group was exposed to a periodic stimulus. Thus, the nature of the Random (sham) stimulus controlled for potential non-specific effects of the flickering stimulus, such as modulation of sensory organs, arousal, or behavioral responses to flickering stimuli. The spiking activity of 1953 neurons and local field potentials (LFP) for 16 animals over 30 sessions were recorded from dorsal hippocampal areas CA1 and CA3 with silicon electrodes as animals performed the VR task or were exposed to flicker (**Figure 1D-F**). This task is relatively simple with a high percentage of correct trials and is likely not very sensitive to memory impairment or enhancement (**Supplementary Figure 1J**). Importantly, following flicker exposure, animals showed similar speed and licking behaviors throughout the track, allowing us to compare neural activity responses directly between stimulation groups without the confounding effects of behavioral differences between groups (**Supplementary Figure 1F-I**).

During slow gamma (25-55 Hz), CA3 and CA1 are more coherent, and therefore CA3 inputs are more likely to influence CA1 activity, supporting memory processes including encoding and retrieval (Bieri et al., 2014; Colgin et al., 2009; Dvorak et al., 2018; Trimper et al., 2014, 2017). Because the frequency of our flicker stimulus is within the slow gamma range, we wondered if 40 Hz flicker might engage similar processes as endogenous slow gamma, resulting in coordinated activity between CA3 and CA1 in the slow gamma range. Thus, we first asked if 40 Hz flicker was affecting communication between these two subregions. We hypothesized that CA3-CA1 coordination would be elevated during exposure to 40 Hz flicker, while the stimulus is having direct effects on the circuit, but that this increase would not persist beyond periods of flicker exposure. To assess coordinated activity across hippocampal subregions, we measured coupling between spike timing and the phase of the local field potential where increased spike-LFP coupling indicates stronger synchronization of the two regions. To quantify CA3-CA1 coupling, we calculated the strength of the interactions between CA3 spiking and CA1 LFP using pair-wise phase consistency (PPC) (**Figure 2A-B)**. PPC is a measure of spike-phase coupling that is not biased by observation metrics such the number of spikes recorded. Positive PPC values indicate that the cell is modulated by a particular LFP phase, a value of 0 indicates there is uniform phase modulation, and negative PPC values indicate anti-phasic modulation (multiple modulated phases that are greater than 90 degrees apart) (Vinck et al., 2012). We specifically filtered out oscillations at 40 Hz to examine coherence that was not directly due to stimulus modulation. We examined PPC between CA1 LFP and both interneurons and pyramidal cells in CA3 because slow gamma activity is thought to be generated and driven by CA3 and depend on both local pyramidal cells and interneurons (Csicsvari et al., 2003). In contrast to our hypothesis, we did not detect a significant change in pyramidal cell CA3-CA1 PPC in the slow gamma band during flicker stimulation or after prolonged exposure to 40 Hz flicker (**Figure 2C, left)**. We found a trend of elevated interneuron CA3-CA1 PPC during flicker after prolonged exposure, but it was not significantly different from Random (**Figure 2D**, left). In contrast, we discovered that CA3-CA1 PPC of both of pyramidal cells and interneurons was significantly increased during VR behavior after prolonged 40 Hz flicker (**Figure 2C-D, right)**. We found on average a 184% increase in CA3 pyramidal cell-CA1 LFP PPC and 130% increase in CA3 interneuron-CA1 LFP PPC following exposure to 40 Hz versus Random flicker (**Figure 2C-D, right)**. While we were primarily concerned with the coupling of CA3 units to CA1 LFP, we observed that these cells also show phase preferences to local CA3 LFP, as expected based on prior work (**Supplementary Figure 4A-B**). Consistent with increased spike-phase coupling, during navigation we found enhanced CA3-CA1 LFP phase synchrony within the slow gamma band following 40 Hz flicker compared to Random (**Supplementary Figure 3A**). These post exposure changes may be due to 40 Hz flicker-induced plasticity in the CA3-CA1 circuit shown *in vitro* in prior studies (Zheng et al., 2020). Our observation of increased CA3-CA1 coupling only during VR behavior may reflect the importance of CA3 during these periods for spatial processing and temporal codes. These results show that prolonged 40 Hz flicker leads to increased functional coupling between CA3 and CA1 in the slow gamma band during navigation.

**Figure 2.**
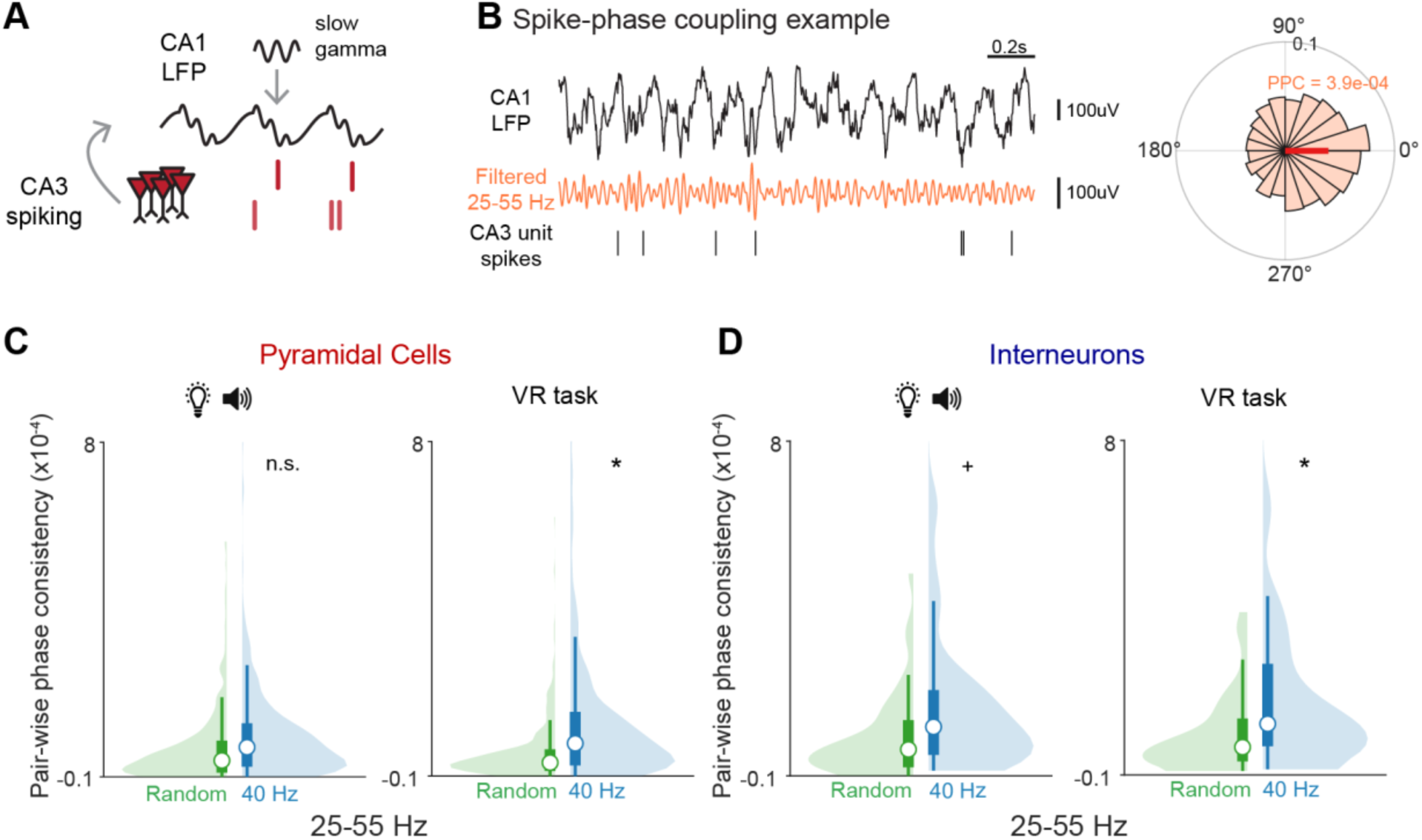
CA3-CA1 spike-phase coupling is increased after 40 Hz flicker. **A.** Pairwise-phase consistency (PPC) quantifies the spike-phase relationship between CA3 single unit spiking and CA1 LFP during slow gamma, which is important for coupling CA3 and CA1. **B.** Slow gamma (25-55 Hz) spike-phase coupling example. *Left*, CA1 LFP (top, black), LFP filtered from 25-55 Hz (middle, orange), example CA3 single unit spikes (bottom, black). *Right*, polar plot showing the spike-phase preference of the example cell to the example CA1 LFP. The polar histogram shows the spiking probability (0 to 0.1) as a function of LFP phase (degrees), and the preferred phase is indicated by the dark orange line. The calculated PPC value is show in orange. Higher positive values indicate stronger synchronization of the two regions. PPC values for endogenous activity are are typically low and are lower for frequencies faster than theta. **C.** 40 Hz flicker results in higher slow gamma spike-field PPC between spiking from putative CA3 pyramidal cells and CA1 LFP during VR behavior periods (right) than Random. There was no significant change during flicker stimulation periods (left). *Left,* flicker stimulation periods, Random, n = 320 pyramidal cells from 8 mice; 40 Hz, n = 387 pyramidal cells from 8 mice; P=0.433, n.s. linear mixed-effects model (LME). *Right*, VR task periods, Random, n = 347 pyramidal cells from 8 mice; 40 Hz, n = 335 pyramidal cells from 8 mice; P=0.024*, LME. See Supplementary Table 2 for statistical details. Violin plots throughout include a histogram (lighter shaded area) and box plot indicating median (white circle), first and third quartiles (dark box), and whiskers (thin lines). **D.** 40Hz flicker results in higher slow gamma spike-field PPC between spiking from putative CA3 interneurons and CA1 LFP during VR behavior. There was a similar trend during flicker periods. *Left*, flicker stimulation periods, Random, n = 46 interneurons from 8 mice; 40 Hz, n = 50 interneurons from 8 mice; P=0.097+, LME. *Right*, VR task periods, Random, n = 49 interneurons from 8 mice; 40 Hz, n = 45 interneurons from 8 mice; P=0.021*, LME.

CA3-CA1 communication is modulated by CA1 theta oscillations in that slow gamma is coupled to CA1 most strongly at certain phases of theta (**Figure 2A**) (Belluscio et al., 2012; Colgin et al., 2009). Thus, we also examined PPC values in the theta (6-10 Hz) band (**Supplementary Figure 4C-D)**. To determine if 40 Hz flicker affected coupling in the theta, slow gamma, and medium gamma bands or only in a subset of frequencies, we constructed a linear mixed model and found a significant interaction between flicker condition and LFP frequency band for CA3-CA1 PPC in both pyramidal cells and interneurons (LME, pyramidal cells P < 2e-16; interneurons, P=0.0056; see **Methods** for details). Examining coupling in the theta band, we found that after 40 Hz flicker, pyramidal cell and interneuron CA3-CA1 PPC did not significantly change. Because CA3 and CA1 are coupled by slow gamma activity but not by other frequencies of gamma, we hypothesized that PPC in other frequency bands of gamma would not be affected. We examined pyramidal cell and interneuron PPC in the medium gamma band (60-100 Hz) (**Supplementary Figure 4E-F)** and found that CA3-CA1 PPC was not changed following 40 Hz flicker. In summary, exposure to 40 Hz flicker leads to slow gamma specific increases in CA3-CA1 spike-phase coupling and phase-phase coupling, indicating stronger coordinated activity which is important for interregional communication.

### 40 Hz flicker increases prospective coding during theta sequences

Because 40 Hz flicker increased CA3-CA1 functional coupling, we next asked if flicker enhances hippocampal codes that depend on CA3. During theta oscillations, spatial locations behind, local to, and in front of the animal’s actual location are represented as a function of theta phase, resulting in theta sequences (Dragoi & Buzsaki, 2006; Middleton & McHugh, 2016; Skaggs et al., 1996). These theta sequences integrate information about past, current, and future spatial position over short time scales for memory processes. Because prior work has shown that theta sequences are dependent on input from CA3, our observations of increased CA3-CA1 coupling after 40 Hz flicker led us to hypothesize that 40 Hz flicker would increase the fidelity of theta sequences during behavior (Middleton & McHugh, 2016). To examine the effects of stimulation on theta sequences, we decoded CA3 and CA1 place cell spiking during theta cycles while the animal was running through the virtual track. We identified significant theta sequences representing positions behind, local to, and in front of the animal’s current position (Farooq & Dragoi, 2019; Middleton & McHugh, 2016). Contrary to our hypothesis, we found that overall theta sequence strength was similar between 40 Hz and Random flicker groups (**Supplementary Figure 6G)**. As CA3 inputs to CA1 are known to promote coding of future locations, we next asked if the types of spatial information being represented during these sequences were different. Because it increases CA3-CA1 coordination, we hypothesized that 40 Hz flicker would increase the coding of future locations (Bieri et al., 2014). To determine the relative amount of forward versus behind information being represented during these sequences, we computed a prospective coding ratio. The prospective coding ratio measures the relative representation of prospective versus retrospective locations as a function of theta phase, with positive values indicating locations in front of the animal’s current position are more strongly represented, and negative values indicating that locations behind the animal are more strongly represented (**Figure 3C**). Surprisingly, we found significantly higher prospective coding ratios after 40 Hz flicker than Random, showing more coding of future positions during theta (**Figure 3D-E**). Following 40 Hz flicker the prospective coding ratio was positive, signifying more prospective coding, while after random flicker the ratio was negative, indicating more retrospective coding. Furthermore, there was a more than two-fold increase in prospective coding on average after 40 Hz than after Random. This 40 Hz flicker-induced increase in prospective coding ratio occurred in the approach to and entry into the reward zones of the track, areas that are especially task relevant. In the 40 Hz group, prospective coding was significantly higher in the reward-related areas of the track as compared to an equally sized control area located half the distance to the next reward location (**Supplementary Figure 6J**). Thus, 40 Hz flicker increased representations of future positions during theta.

**Figure 3.**
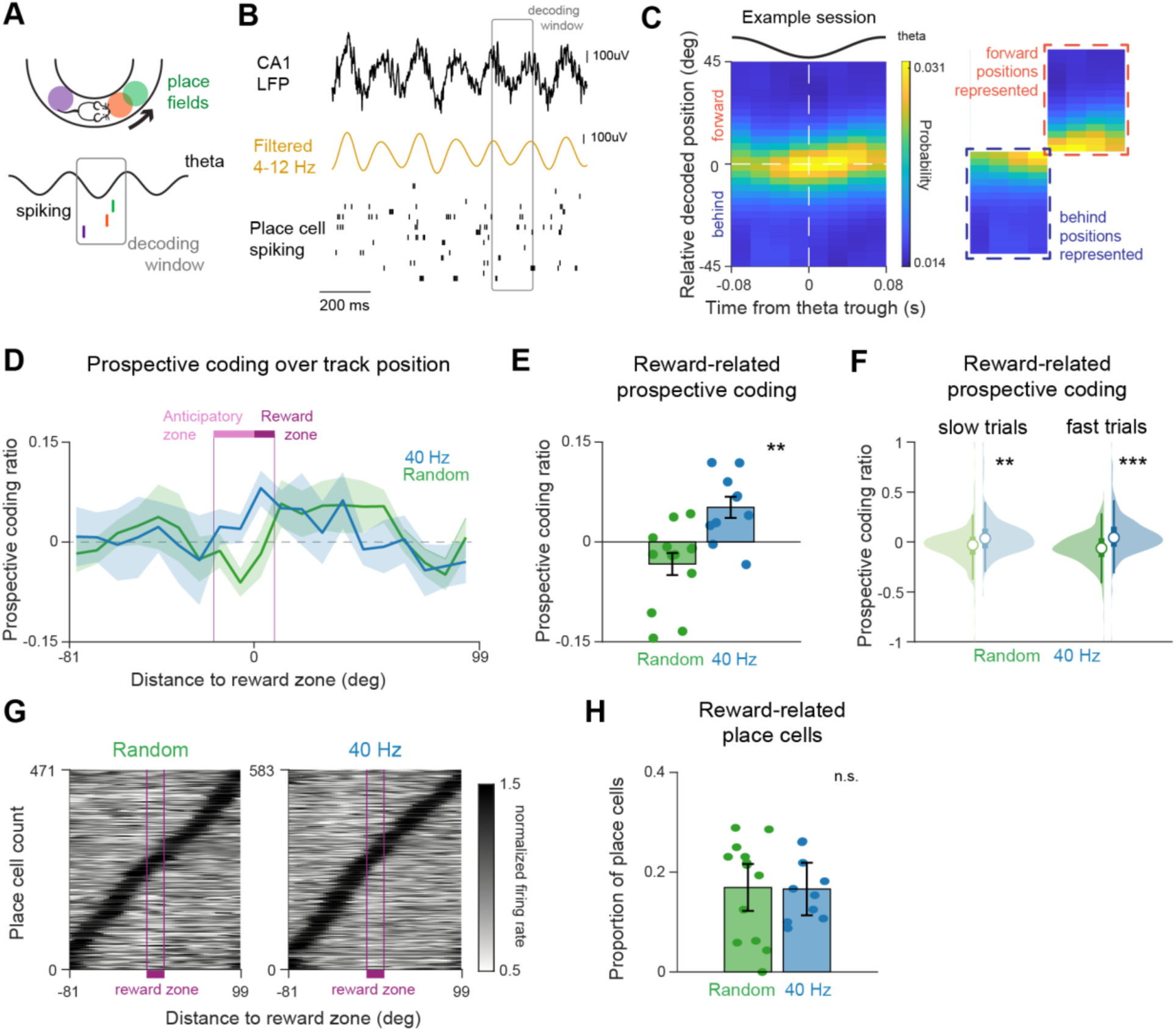
40 Hz flicker increases prospective coding during theta. **A.** Illustration of spiking as a function of theta phase. Descending phases of theta tend to represent positions behind the animal’s current location (purple), while ascending phases of theta represent positions in front of the current position (orange and green). The grey box indicates the time window used in the decoding relative to theta phase. **B.** Example of CA1 LFP and place cell spiking activity used to calculate theta sequences. CA1 LFP trace (top, black), LFP filtered from 4-12 Hz (theta band; middle, dark yellow), and place cell spiking activity (bottom, black lines indicate spikes) with each row a different neuron. The grey box indicates the time window used in the decoding in **C** relative to theta phase. **C.** *Left*, an example recording session illustrates an average theta sequence with a higher probability (more yellow) of decoding positions in front of the animal (positive relative decoded position) on the rising phase of theta and a higher probability of decoding positions behind the animal (negative relative decoded position) on the falling phase. *Right,* the decoded data is divided into quadrants, and the forward quadrant representing positions in front of the animal (within orange dashed box, top right) and the behind quadrant representing positions behind the animal (within dark blue dashed box, bottom left quadrant) are isolated. A prospective coding ratio is defined comparing the spatial probabilities in the forward versus behind quadrants (see Methods). **D.** Average prospective coding ratio value over position centered around the reward zones after 8 days of 1hr/day of 40 Hz (blue) or Random (green) flicker. In animals exposed to 40 Hz flicker, prospective coding increased as animals approached (anticipatory zone, light pink box) and entered the reward zone (dark pink box). The prospective coding ratio ranges from 1, indicating that only forward positions are represented, to −1, indicating only behind positions are represented with 0 indicating equal representation of forward and behind positions. Mean ± SEM across recording days. **E.** Prospective coding ratios after 8 days of 1hr/day of 40 Hz (blue) were significantly higher than after Random (green) flicker for the theta cycles occurring in the anticipatory and reward (reward-related) zones of the track, as indicated by the pink bar in **D.** One dot indicates the average prospective coding ratio in the reward-related zones on one recording day. Random, n = 13 days from 8 mice; 40 Hz, n = 10 recording days from 6 mice; P = 0.003**, linear mixed-effects model (LME). See Supplementary Table 2 for statistical details. Mean ± SEM across recording days. **F.** Prospective coding ratios in reward-related zones were significantly higher after 40 Hz (blue) than Random (green) flicker for both trials with low speed (left) and high speed (right) after 8 days of 1hr/day flicker. Low speed, Random, n = 480 trials from 8 mice; 40 Hz n = 389 trials from 6 mice; p=0.005**, LME. High speed, Random, n = 454 trials from 8 mice; 40 Hz, n = 418 trials from 6 mice; P=0.0007***, LME. **G.** Normalized firing rate maps for place cells recorded from animals exposed to Random (left) or 40 Hz (right) stimulation as a function of distance to the reward zone. Cells are ordered by peak firing position in the track. Reward zones are indicated by pink lines and boxes along the x-axis. Random, n = 471 place cells from 8 mice; 40 Hz, n = 583 place cells from 8 mice. **H.** Percentage of place cells with fields located in the anticipatory and reward zones of the track did not differ significantly after 8 days of 1hr/day of 40 Hz (blue) or Random (green) flicker. Random, n = 13 days from 8 mice; 40 Hz, n = 10 recording days from 6 mice; P=0.783, n.s. LME. Mean ± SEM across recording days.

Running speed has been shown to affect coding during theta sequences, therefore we examined whether velocity was driving differences in prospective coding between groups (Gupta et al., 2012; Maurer et al., 2012; Parra-Barrero et al., 2021). Overall, 40 Hz and Random flicker animals displayed similar running speed during the approach and entry into the reward zone (**Supplementary Figure 1)**. We further separated trials into fast and slow speed trials within each group (see Methods) and found that trial speeds in these groups were similar between 40 Hz and Random groups. We then examined the prospective coding ratio in the anticipatory and reward zones between the 40 Hz and Random groups in these trial speed subsets. Prospective coding ratio in these reward-related areas after 40 Hz flicker was significantly higher, positive, and over two times greater than Random on average on slow and fast trials. These results show that the observed differences in prospective coding are not due to speed differences between groups (**Figure 3F**).

We then found that these differences in prospective coding are not explained by differences in place cell representations of the approach and entry into the reward zones. Prior work suggests that CA3 is important for supporting temporal codes or the timing of spikes, but not rate coding or the number of spikes of pyramidal cells in hippocampus (Middleton & McHugh, 2016). Thus, we hypothesized that following 40 Hz flicker, place cell properties would remain unchanged. In agreement with this hypothesis, we found that place cells’ spatial and rate coding and the proportions of place cells representing areas of the track leading up to and inside of the reward zone did not differ significantly in the 40 Hz and Random flicker groups (**Figure 3G-H, Supplementary Figure 6A-F)**. These results also indicate that our finding of increased prospective coding during theta is not due to differences in the representation of the rewarded areas of the track between groups. In summary, we found that 40 Hz flicker increased prospective coding during theta, while the spatial properties of place cells and their firing fields remained unchanged. These results show that 40 Hz flicker increases information about future place positions within temporal codes, consistent with increased CA3-CA1 coordination.

### 40 Hz flicker increases prospective coding during sharp-wave ripples

SWRs are essential for spatial memory and impaired in models of AD, and we therefore wondered how 40 Hz flicker affects SWRs. SWR reactivation is dependent on CA3 input which also promotes prospective coding in the hippocampus (Nakashiba et al., 2009). Therefore, we next asked if there were increases in prospective coding during SWRs, where sequences of place cells representing spatial trajectories extending farther from the animal are reactivated on the compressed timescales required for plasticity. Because we found 40 Hz flicker increased CA3-CA1 coupling, we hypothesized that 40 Hz flicker would enhance prospective coding during SWRs. We decoded estimated position content from SWR events using a Bayesian decoder and non-overlapping time bins (**Figure 4A-B**, see Methods). Because SWRs occur during pauses in behavior, we restricted our decoding analysis to SWRs occurring in the reward zone of the track where animals consistently paused while consuming reward. We defined a prospective coding ratio to determine whether the SWR more strongly represented positions forward (ratio values > 0) or behind (ratio values < 0) the animal’s current location (see Methods). We found that following 40 Hz flicker, SWRs had significantly higher positive prospective coding ratio values than following Random flicker, indicating more prospective coding (**Figure 4C, 4D**). The prospective coding ratio increased by over 3-fold on average following 40 Hz flicker versus Random (**Figure 4C, 4D**). To determine the magnitude of prospective coding during SWR events, we applied sequence-less decoding to all SWR events, decoding spiking during the SWR in one time bin to yield a single spatial probability estimate and determined the difference between the animal’s actual location during the SWR and the maximum decoded position. After Random flicker, the relative positions represented were distributed across the track. In contrast, following 40 Hz flicker, SWRs tended to represent positions in front of the animal, mostly those at least 18 degrees (1 track zone) away from the animal’s current position (**Figure 4D**). Furthermore, the fraction of SWRs that preferentially coded for future position was on average 57% higher following 40 Hz flicker than Random (**Figure 4E**). These results show that 40 Hz flicker resulted in higher SWR prospective coding.

**Figure 4.**
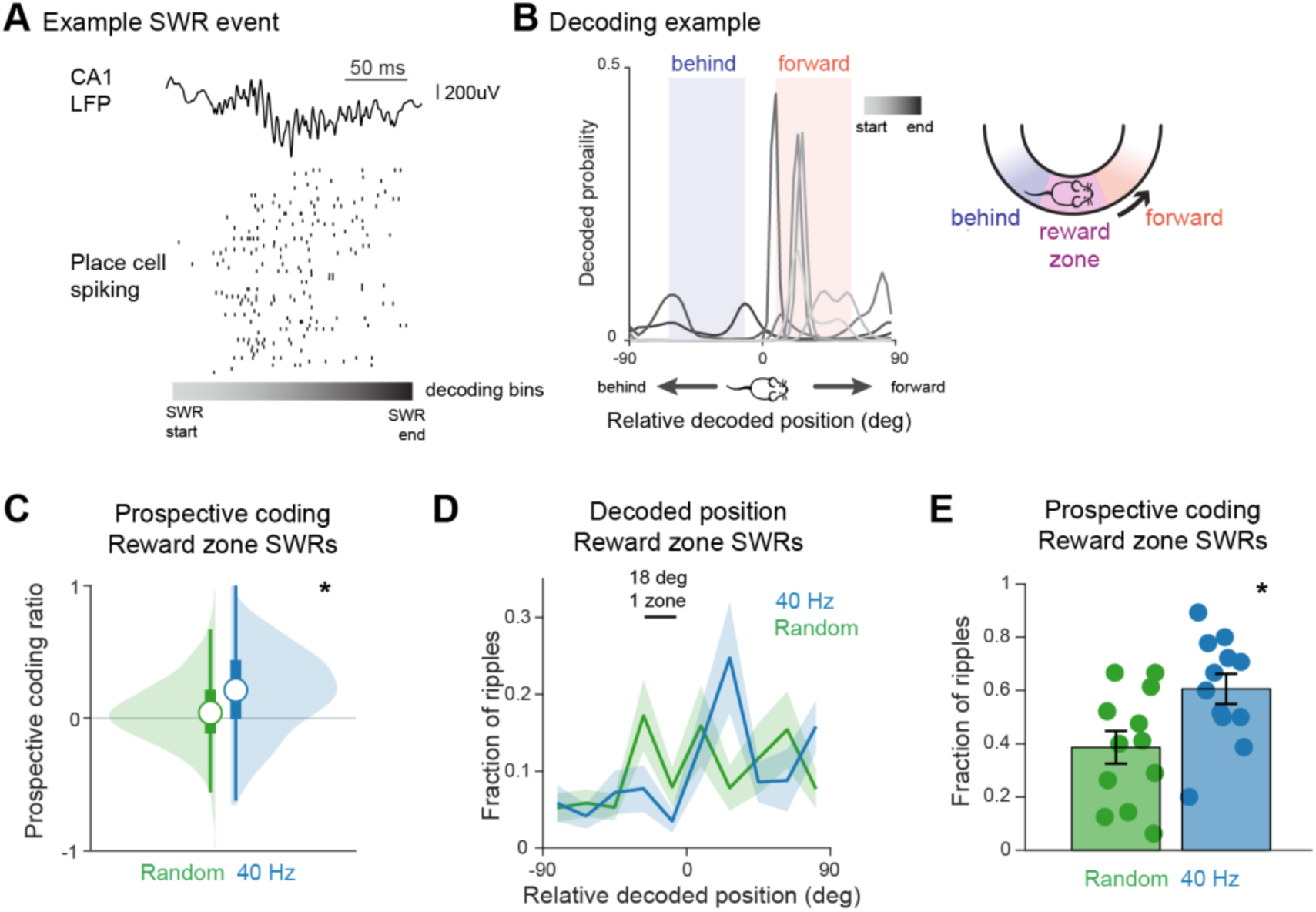
40 Hz flicker increases prospective coding during sharp-wave SWRs. **A.** Example sharp-wave ripple event. *Top*, CA1 LFP. *Bottom*, place cell spiking. Each row corresponds to the spiking activity of one place cell. Place cells are sorted by peak firing rate location in the track. The grey scale heat map below the spiking activity corresponds to the decoding bins shown in **B** from ripple start (light grey) to end (dark grey). **B.** Decoding example for the ripple event shown in **A** shows stronger decoding of positions in front of the animal. *Left*, spatial probabilities of decoding a position in the track relative to the animal’s current position (at zero) during each time bin of the ripple event. Decoded spatial probabilities from temporal bins early (lighter shades of grey) and in the middle (medium shades) of the ripple event more strongly represent forward positions while the late bins in the ripple event (darker shades of grey), represent positions behind the animal in this example. Spatial probabilities representing positions in front of the animal (orange shaded area) were summed and compared to spatial probabilities representing positions behind the animal (dark blue shaded area) to compute a prospective coding ratio value. *Right*, schematic indicating forward (orange) and behind (dark blue) positions relative to the reward zone (pink). **C.** Prospective coding ratio values for all SWRs occurring in the reward zone of the track were higher after 8 days of 1hr/day of 40 Hz (blue) than Random (green) flicker, indicating stronger decoding of forward positions. A prospective coding ratio of 1 indicates the ripple represented only positions forward of the animal, while −1 indicates the ripple represented only positions behind the animal. Random, n = 507 SWRs from 7 mice; 40 Hz, n = 378 SWRs from 7 mice; P = 0.047*, LME. See Supplementary Table 2 for statistical details. **D.** Relative decoded position for all SWRs occurring in the reward zone of the track after 8 days of 1hr/day of 40 Hz (top) or Random (bottom) flicker shows more decoding of positions forward of the animal (> 0) after 40 Hz flicker, with a majority of SWRs representing positions 1-2 zones (18-36 degrees) in front of the animal’s current position. The relative decoded position is the difference between the decoded position during SWRs and the actual position when the ripple occurred, with positive values indicating decoded positions in front of the animal and negative values indicating decoded position behind the animal. Mean ± SEM across recording days, Random, n = 12 days from 7 mice; 40 Hz, n = 12 days from 7 mice. **E.** A higher proportion of SWRs had positive prospective coding after 8 days of 1h/day of 40 Hz (blue) than Random (green) flicker. SWRs with prospective coding values greater than 0 contain more prospective than retrospective content; ratio values greater than 0.1 were deemed prospective coding SWRs to exclude SWRs primarily representing the current position. Random, n = 12 days from 7 mice; 40 Hz, n = 12 days from 7 mice; P = 0.033*, LME. Mean ± SEM across recording days.

Changes in prospective coding during SWRs led us to ask if there were underlying improvements in SWR properties. SWR properties including abundance, size, and duration have been previously found to be deficient in mouse models of AD (Gillespie et al., 2016; Jones et al., 2019; Prince et al., 2021). The duration of SWR events has been associated with improved memory, and SWRs have been shown to be shorter in 5XFAD mice (Fernández-Ruiz et al., 2019; Jones et al., 2019; Prince et al., 2021). We examined the duration of SWR events, the abundance of SWRs during these non-theta periods, and SWR size and found that these properties were not significantly changed following 40 Hz flicker (**Supplementary Figure 7D-F**). Activation and co-activation of place cell pairs during SWRs have been implicated in spatial memory task performance and are deficient in 5XFAD mice (Prince et al., 2021; Singer et al., 2013). We found no significant differences between flicker groups in the likelihood of place-cell pairs to be active or co-active together during SWRs (**Supplementary Figure 7G-H)**. Our results show that 40 Hz flicker increased prospective coding during SWRs, despite a lack of significant changes in SWR abundance, duration, size, and co-activation.

### Prospective coding during theta sequences is increased during efficient task behavior

Having identified that 40 Hz flicker increases prospective coding, we evaluated how this prospective coding relates to task performance. Prospective coding is thought to dynamically increase to enable animals to plan farther ahead when moving more quickly. Indeed, prior work has shown that theta sequences are longer and code farther in front of the animal when animals run faster (Gupta et al., 2012; Parra-Barrero et al., 2021). As such, we wondered if more prospective coding was correlated with faster and more efficient behavioral performance in our task in 5XFAD mice, similar to prior work in healthy animals. We found that prospective coding across the track was significantly higher on fast trials than slow trials, in both animals exposed to 40 Hz and Random stimulation (**Figure 5D-E**). This increase in prospective coding on fast trials occurred immediately after the reward zone, where animals started the next trial following reward delivery. Quantifying the prospective coding ratio for theta cycles occurring in this post-reward region of the track when animals initiate their run to the next reward zone, we found that in both stimulation groups fast trials had significantly more prospective coding than slow trials (**Figure 5E**). In our task, running quickly results in the animal receiving more rewards in a shorter period and thus is an advantageous behavioral strategy for maximizing receipt of rewards (**Figure 5F**). These results are consistent with prior work in healthy animals showing that prospective coding increases dynamically during efficient goal-directed navigation. As licking is crucial to successful performance of this task, we asked if prospective coding was correlated with licking behavior. After 40 Hz flicker, prospective coding ratio was significantly higher on trials with higher anticipatory licking (**Supplementary Figure 8A)**. These results show that more prospective coding during theta is correlated with high-speed running behavior, a more efficient approach to accumulate rewards in this task. Furthermore, after 40 Hz flicker, prospective coding is higher when animals lick more in anticipation of reward.

**Figure 5.**
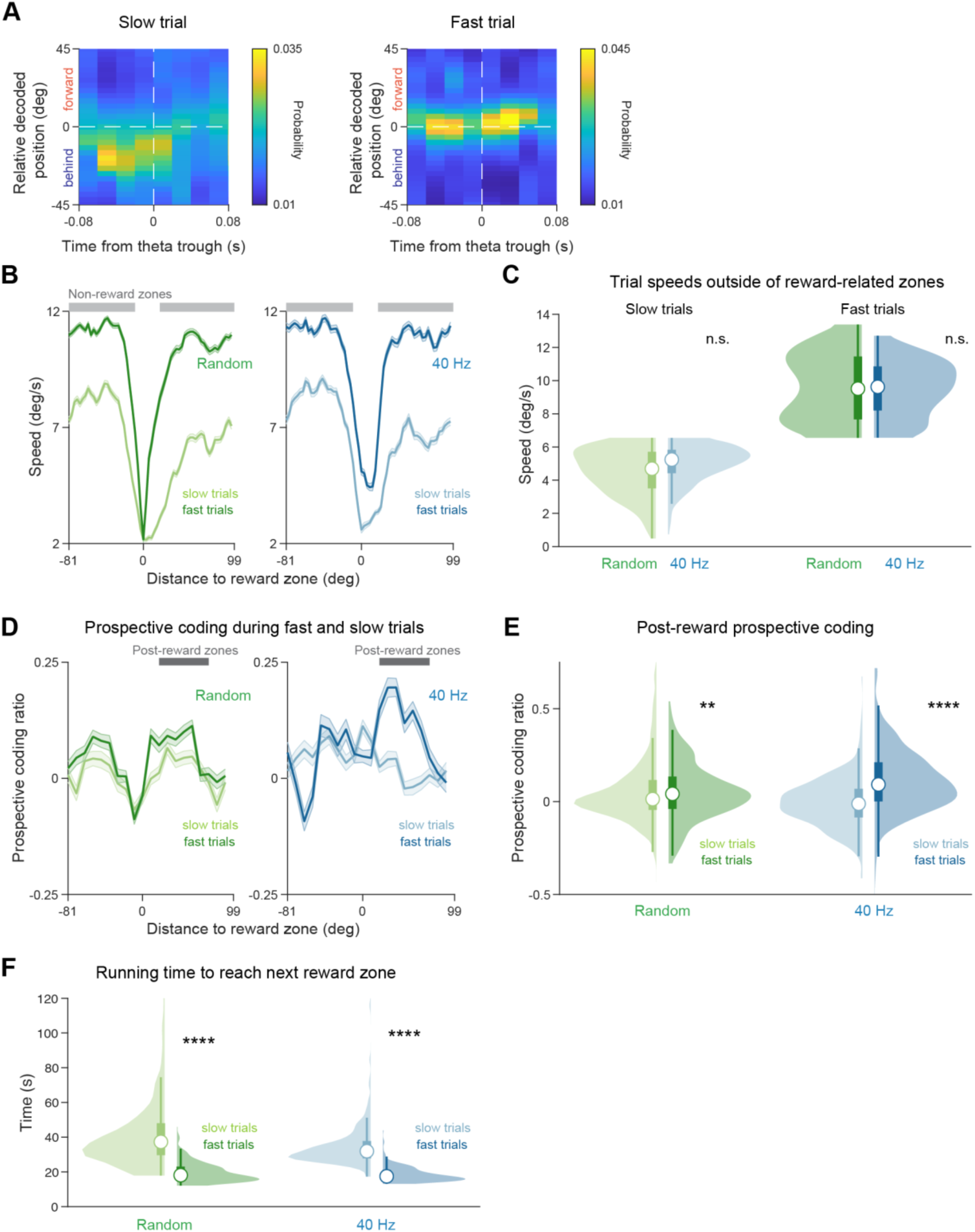
Prospective coding during theta is higher during efficient task behavior. **A.** Example average theta sequence computed from theta cycles occurring after the reward zone during a slow trial (left) with stronger decoding of position behind the animal and a fast trial (right) with stronger decoding of forward positions from one animal. **B.** Average speed across the track for trials in the top half (fast trials, dark colors) and bottom half (slow trials, light colors) of trial speeds outside of the rewarded areas of the track after 8 days of 1hr/day of Random (left, green) or 40 Hz (right, blue) flicker. Included trials: random, slow n = 483 trials, fast = 454 trials, from 8 mice; 40 Hz, slow n = 389 trials, fast n = 418 trials, from 6 mice. Light grey bars indicate non-rewarded areas of the track. Speed in the non-rewarded areas was used to separate trials into fast and slow trials (grey bars indicate non-rewarded areas). Mean ± SEM across trials. **C.** Distributions of trial speeds in the bottom half (slow, left) or top half (fast, right) of all trials after 8 days of 1hr/day of Random (green) or 40 Hz (blue) flicker. Speeds in slow trials and fast trials did not differ significantly between stimulation groups. Slow trials, Random, n = 483 trials from 8 mice; 40 Hz n = 389 trials from 6 mice; P=0.214, n.s. LME. Fast trials, Random, n = 454 trials from 8 mice; 40 Hz, n = 418 trials from 6 mice; P=0.993, n.s. LME. **D.** Per-trial average prospective coding ratio value across the track after 8 days 1hr/day of Random (left, green) or 40 Hz (right, blue) flicker. During high-speed trials, prospective coding was higher than low-speed trials in both stimulation groups in the post-reward zones of the track (dark grey horizontal bar), where the animal begins running to the next rewarded zone. Random, slow n = 483 trials, fast = 454 trials, from 8 mice; 40 Hz, slow n = 389 trials, fast n = 418 trials, from 6 mice. Mean ± SEM across trials. **E.** Prospective coding ratio was higher on fast than slow trials in both stimulation groups. Ratios were calculated per-trial for theta cycles in the post-reward zones of the track, indicated by the dark grey rectangle in **D.** Random, slow n = 483 trials, fast = 454 trials, from 8 mice; P=0.0086**, LME. 40 Hz, slow n = 389 trials, fast n = 418 trials, from 6 mice; P=1.1e-9****, LME. **F.** Less time was required to reach the next reward zone on fast than slow trials in both flicker groups. Random, slow n = 475 trials, fast = 454 trials, from 8 mice; P<2.2e-16**** LME. 40 Hz, slow n = 386 trials, fast n = 417 trials, from 6 mice; P<2.2e-16****, LME.

To determine if higher prospective coding ratio during fast trials was a consequence of faster motor behavior, we examined prospective coding during trials where the mice were running through the track but failed to stop and lick at any location in the track (“unengaged trials”). On these trials, mice ran significantly faster than on trials where animals licked for a reward in the reward zone (“engaged trials”, **Supplementary Figure 8B**). We hypothesized that if higher prospective coding ratios were purely due to running speed, unengaged trials would have higher prospective coding ratios regardless of task performance. In contrast, we find that on unengaged trials prospective coding ratio is significantly lower than during correct trials, even though animals are running faster (**Supplementary Figure 8C**). When we consider 40 Hz and Random flicker groups separately, we find prospective coding ratio is significantly higher on correct trials than unengaged trials in animals that underwent 40 Hz flicker and no significant difference between trial types in the Random group (**Supplementary Figure 8D-E**). Thus, higher prospective coding on fast trials is not due solely to faster running speed in our task.

The prospective coding ratio during theta represents nearby prospective positions, corresponding to one or two zones ahead of the animal. Because the next reward zone is the animal’s next goal and a salient part of the track, we asked if trials ever included coding for the next goal location, or unoccupied reward zone. After 40 Hz flicker, we found that many trials contained representation of the next goal location, and this decoding was specific to the rewarded area (**Supplementary Figure 9A**). We find that these trials have similar types of behavior performance, though after 40 Hz flicker slow trials were more likely to contain decoding of the unoccupied reward zone than fast trials (**Supplementary Figure 9B-E**). We also examined representation of the unoccupied goal location during SWRs and found that about half of SWRs contain representation of the unoccupied reward location **(Supplementary Figure 10A**). Behavioral performance did not differ significantly between trials with and without unoccupied reward zone representations during SWRs (**Supplementary Figure 10B-D**).

Thus, we show that 40 Hz flicker exposure leads to increased prospective coding that is associated with efficient and engaged task performance, including faster running and more anticipatory licking, strategies that lead to more rapid rewards. Prospective coding increases after 40Hz flicker are not purely a consequence of fast running speed since animals ran faster on incorrect “unengaged” trials than correct trial.

## Discussion

Although gamma sensory flicker is being tested as a potential intervention in AD, how this stimulation acts on memory processes has been unclear. Here we addressed this gap by investigating the effects of 40 Hz flicker on neural activity essential for memory in hippocampus. We observed that 8 days of exposure to 40 Hz flicker compared increased functional coupling between hippocampal subregions CA3 and CA1 increased during navigation, specifically coupling within the slow gamma band, compared to Random (sham) flicker. CA3-CA1 interactions play a central role in generating sequences that integrate information about past, present, and future experiences and CA3 inputs to CA1 are important for prospective coding (Bieri et al., 2014; Middleton & McHugh, 2016). Therefore, we tested the hypothesis that 40 Hz flicker increases such hippocampal sequences. We found that in reward-related areas of the track, 40 Hz flicker increased prospective coding during both theta sequences and SWRs, indicating more representation of future positions during network states associated with both active running behavior and pauses in behavior. This observed increase in prospective coding was independent of significant changes in place field and SWR properties. Finally, we found that increases in prospective coding are correlated with more efficient behavioral strategies. Prospective coding during theta sequences was increased during more efficient trials with higher running speeds. Prospective coding was higher in animals exposed to 40 Hz than those exposed to Random flicker in both high and low speed trials, showing that while the magnitude of the prospective coding ratio varies with speed, differences in speed did not explain 40 Hz flicker-induced increases in prospective coding. After 40 Hz flicker, prospective coding is higher when animals lick more in anticipation of reward. Together, these results show that chronic 40 Hz flicker enhances specific aspects of hippocampal activity that are important for memory encoding and retrieval, namely CA3-CA1 interactions and prospective coding during theta and SWRs.

40 Hz flicker induced improvements in hippocampal subregion coupling as compared to Random flicker. Specifically, CA3-CA1 coupling increased during VR navigation after but not during 40 Hz flicker exposure. This finding suggests that 40 Hz flicker leads to stronger inputs from CA3 to CA1. These results do not mean that 40 Hz flicker is entraining endogenous slow gamma such that the endogenous oscillation is synchronized to the stimulus (Soula et al., 2023). Rather, the impact of stimulation in the circuit results in increased CA3-CA1 coupling in the slow gamma band after stimulation is ceased. It is important to note that some studies have used the word “entrainment” in different ways. Several early studies on gamma sensory stimulation used “entrainment” to describe how an external 40 Hz sensory stimulus causes an increase in 40 Hz frequency neural activity in the brain. More recently, Soula *et al*. used the term “entrainment” to mean an external rhythmic stimulus synchronizing internal endogenous oscillations to the stimulus. Endogenous oscillations are of interest because of their functional roles in circuit computations; slow gamma oscillations are thought to facilitate CA3-CA1 interactions. Thus, we asked how 40 Hz flicker affects such circuit computations, specifically, coupling between CA3 and CA1 that occurs during endogenous slow gamma activity. We theorized that 40 Hz flicker could affect this key function of slow gamma oscillations regardless of whether endogenous slow gamma becomes significantly synchronized to the stimulus. 40 Hz flicker may enhance CA3-CA1 coupling by recruiting intrinsic resonant properties in the circuit. Because CA3 excitatory-inhibitory connections generate slow gamma activity, they might preferentially be engaged by 40 Hz stimuli (Csicsvari et al., 2003). While 40 Hz sensory flicker is quite different from endogenous oscillations, part of the rationale for using this stimulation frequency is that it may induce some of the effects of endogenous slow gamma. Indeed, recent work has shown that 40 Hz visual flicker improves CA3-CA1 long-term potentiation (LTP) in a stroke model, which suggests a plasticity mechanism underlying improvements in CA3-CA1 subregion communication (Zheng et al., 2020). However, it remains to be determined if similar plasticity mechanisms are engaged in AD models. We conclude that 40 Hz flicker increases CA3-CA1 coupling during navigation.

CA3-CA1 coordination has functional importance in the hippocampus because CA3 is essential for memory processes, including the generation of neural sequences that integrate information about past, present, and future (Middleton & McHugh, 2016; Nakashiba et al., 2009; Nakazawa et al., 2003). We found that following 40 Hz sensory stimulation, prospecting coding is increased during theta sequences and SWRs. Increases in prospective coding during these sequences could be due to stronger CA3 influences on CA1 or due to stronger sequence generation within CA3. Gamma sensory stimulation has multiple reported effects, including lowering amyloid beta levels, altering microglia and cytokines, mitigating synaptic and neuronal loss, and increasing glymphatic clearance (Adaikkan et al., 2019; Iaccarino et al., 2016; Martorell et al., 2019; Murdock et al., 2024; Prichard et al., 2023). These different effects of gamma sensory flicker may contribute to flicker-induced increases CA3-CA1 coupling and prospective coding. Prior evidence suggests that CA3 is necessary for temporal coding, like spiking sequences, but not rate coding in CA1 (Middleton & McHugh, 2016). In agreement with this prior work, our results show increased prospective coding during theta sequences and SWRs, following increased CA3-CA1 coordination induced by 40 Hz flicker, without changes in spatial rate coding.

As prospective codes represent future information and are thought to be important for planning for upcoming decisions, we examined how prospective coding relates to behavioral performance (Hasz & Redish, 2018; Wikenheiser & Redish, 2015). Consistent with prior work showing that the representation of forward locations increases with higher running speeds, we find that prospective coding during theta sequences increased during trials with higher running speeds (Gupta et al., 2012; Maurer et al., 2012; Parra-Barrero et al., 2021). We also found that slow trials were more likely to contain decoding of the unoccupied reward zone than fast trials after 40 Hz flicker. We speculate that these slower trials and distal prospective coding may reflect a more deliberative state of the animal. We examined how prospective coding relates to task engagement. Interestingly, after 40 Hz flicker prospective coding was higher on engaged correct trails and higher when animals licked more in anticipation of reward. Importantly, these findings show prospective coding increases after 40 Hz flicker are not purely a consequence of fast running speed since animals ran faster on incorrect “unengaged” trials than correct trials. After 40 Hz flicker, prospective coding was increased in the areas of the track immediately following the reward zones, during which prospective coding may be advantageous in directing behavior on the next trial. Together these results show that 40 Hz flicker increases prospective coding, and these increases are correlated with more task engagement and efficient and effective task performance.

We find that animals exposed to 40 Hz flicker have a higher proportion of SWR events representing prospective information as compared to animals exposed to Random flicker. We examined multiple properties of SWRs including SWR duration, abundance, and related place cell coactivation, and found that these metrics were unchanged following flicker. The lack of significant changes in SWR abundance agrees with prior work conducted in mice that express *APOE4*, a variant of apolipoprotein that is the primary genetic risk factor for AD. *APOE4* knock-in mice display deficits in SWR abundance and SWR-associated slow gamma. Removal of *APOE4* from GABAergic interneurons rescued SWR-associated slow gamma, but SWR abundance was unchanged (Gillespie et al., 2016). The effects of APOE4 on SWR replay content, like prospective codes, have yet to be determined. Importantly, our findings indicate that the increases in prospective coding during SWRs were not merely due to changes in the underlying abundance of SWR events. These results show that both 40 Hz flicker and removal of *APOE4* from GABAergic interneurons result in improvement in the quality of SWR activity without rescuing SWR abundance.

40 Hz flicker results in increased hippocampal CA3-CA1 coordination and prospective coding in multiple network states, even when behavior is similar between 40 Hz and Random flicker groups following flicker exposure. Prior work showing improvements in memory following 7-days of 40 Hz flicker was conducted in the same mouse model at a similar time point (5XFAD, 6 months) using tasks designed to detect improvements in behavior (Martorell et al., 2019). Yet it remained unclear how sensory modulation of 40 Hz activity in the hippocampus could improve memory processes, as such sensory-evoked activity is not known to or theorized to play a role in memory. Thus, we aimed to understand how 40 Hz sensory stimulation affects neural activity required for learning and memory, a key gap in elucidating the effects of this stimulation. We addressed this question using a simple paradigm designed such that 5XFAD mice could learn to perform the task well, and animals were well trained prior to and throughout the recording experiment. Because hippocampal codes can depend on or vary with behaviors such as running speed, using a task that results in similar behavioral performance across groups is important (Maurer et al., 2012; McNaughton et al., 1983). We were able to directly compare neural activity differences between the two groups and conclude that observed changes were due to flicker condition, not gross behavioral differences. Of course, there may be more subtle behavioral differences that we were unable to detect. While relatively simple, this task enabled measurement of neural activity that has been shown to be important for memory. We focused on hippocampal circuits and specifically CA3 and CA1 subregions because of the wealth of literature on slow gamma in these regions and the extensive research on patterns of activity essential for memory in these circuits (Bieri et al., 2014; Carr et al., 2012; Colgin et al., 2009; Dvorak et al., 2018; Fernández-Ruiz et al., 2019; Girardeau et al., 2009; Jadhav et al., 2012; Montgomery & Buzsáki, 2007; Trimper et al., 2014). However, our findings do not preclude flicker induced changes in cortical neuronal dynamics. We examined CA3-CA1 coherence because naturally occurring increases in CA3-CA1 coherence during encoding predict better memory recall (Trimper et al., 2014, 2017). Furthermore, naturally occurring increases in prospective coding, like those induced by 40 Hz flicker, have been shown to co-occur with endogenous gamma and predict correct spatial memory (Bieri et al., 2014; Dvorak et al., 2018). Thus, the effects of 40 Hz flicker observed in this simple task are analogous to changes in neural activity shown to be important for memory encoding and recall. This study is an essential step to determine which kinds of activity are affected by stimulation and could be contributing to memory enhancement to identify what activity should be disrupted in a causal test.

Understanding first how stimulation affects patterns of activity essential for memory is crucial to design a sophisticated and appropriate test of circuit mechanisms. Further work is needed to causally test if these specific changes account for memory improvement in response to stimulation. From the findings presented here, we hypothesize that CA3 inputs to CA1 during and immediately after stimulation are required for 40Hz flicker’s effects on both prospective coding and memory. This is in line with studies that have shown sensory flicker increases CA3 to CA1 LTP (Zheng et al., 2020).

We find that exposure to 40 Hz audio-visual flicker improves endogenous memory-related neural activity in the hippocampal circuit, specifically resulting in increased CA3-CA1 coordination and enhanced prospective coding during theta and SWRs. These results reveal new insights into how 40 Hz flicker affects neural mechanisms of memory. Based on the role of these patterns of activity in hippocampal function and memory, these findings suggest candidate mechanisms through which 40 Hz flicker acts on network activity to improve cognition.

## Methods

### Animals

All animal work was approved by Georgia Institute of Technology’s Institutional Animal Care and Use Committee. The mouse strain used for this research project, B6SJL-Tg(APPSwFlLon,PSEN1*M146L*L286V)6799Vas/Mmjax, RRID:MMRRC_034840-JAX, was obtained from the Mutant Mouse Resource and Research Center (MMRRC) at The Jackson Laboratory, an NIH-funded strain repository, and was donated to the MMRRC by Robert Vassar, Ph.D., Northwestern University. 5XFAD mice were bred on a C57Bl/6 background in the animal facilities at Georgia Institute of Technology. 6–7-month-old male 5XFAD mice were used for all experiments. Littermates were split between experimental groups when applicable. Mice were single-housed on a reverse 12-hour light/12-hour dark cycle. Behavioral training, electrophysiology experiments, and flicker exposure were performed during the dark cycle. At the start of behavioral training, mice were food-restricted between 85-90% percent of their baseline body weight, and food restriction continued for the duration of the experiment. Water was provided without restriction. Animals were excluded from the experiment if they did not display behaviors sufficient to perform the behavioral task (N = 1 mouse) or had surgical complications (N = 2 mice).

### Headplate implantation and craniotomies

During headplate implantation surgery, mice were anesthetized with isoflourane, a custom stainless steel headplate was fixed using dental cement (C&B Metabond, Parkell), and the target craniotomy site for LFP recordings was marked on the skull (in mm, from bregma: −2.93 anterior/ posterior, +2.1 or −0.98 medial/lateral for targeting CA1; −1.6 anterior/posterior, +/-1.6 medial/lateral for targeting CA3). Following behavioral training, craniotomies (300-500μm in diameter) were made by thinning the skull with a dental drill and then making a hole with a 27-gauge needle. Two sets of craniotomies were performed, the first following behavioral training, and the second set following 8 days of flicker exposure. The hemisphere for the first set of craniotomies was chosen randomly, and the second set of craniotomies was performed on the opposite hemisphere. The craniotomy was sealed with a sterile silicon elastomer except when performing electrophysiology recordings (Kwik-Sil WPI).

### Behavioral training and analysis

The virtual environment was a continuous annular track, with different visual cues on the walls every 18 degrees (“zones”) and was projected onto a cylindrical screen in front of the animal (Aronov & Tank, 2014). Because the full track spans 360 degrees, there were 20 zones in the environment each with a distinct wall pattern. The reward zones were located 180 degrees apart from each other. Head-fixed animals ran on a spherical treadmill composed of an 8-inch polystyrene foam ball floating on air and were exposed to the virtual reality environment the first time they were head-fixed on the treadmill. As animals ran on the treadmill, the translational and rotational velocities were tracked via an optical mouse and converted into movement through the virtual reality environment. In the first phase of behavioral training, animals navigated around the annular track environment and automatically received a sweetened condensed milk reward (1:2 water dilution) when they entered either of the two reward zones. During training, 3 drops of sweetened condensed milk reward were delivered per reward zone. Once animals comfortably ran through the track and began demonstrating anticipatory licking before the reward zone, the animals were transitioned to the second phase of behavioral training where they were required to lick in the reward zones to receive the reward. Licks were detected using a photo-interrupter placed in front of the animal’s mouth around the reward spout. Electrophysiology recordings commenced once animals were able to perform the second phase of the behavioral task reliably. During recordings, the amount of reward delivered in the virtual reality track was increased to 4 drops of sweetened condensed milk to encourage long durations of pausing during the task, as sharp-wave ripples occur during pauses in behavior. As animals performed the VR task, behavioral data was collected, including translational and rotational velocity, licking, position, and reward delivery. A moving average was used to smooth position and velocity data in MATLAB. Behavioral data was segmented into trials in which each trial was either one full lap (360 degrees) around the annular track in which the animal passed through two reward zones, or one half lap (180 degrees) in which the animal passed through one reward zone. Licking activity was quantified as the fraction of licks in each position bin over a trial, and speed was quantified as degrees traveled in the track per second. Behavior was similar between the two halves of the track, and the data was combined as a function of distance to the reward zone for most analyses. Correct, engaged trials were trials on which an animal licked in the reward zone and thus received a reward. On incorrect, unengaged trials, animals ran through the track but did not lick in any zones of the track, including the reward zone, and therefore did not receive a reward.

### Electrophysiology recordings

Recording sessions lasted a maximum of 5 hours. Animals were free to run or rest on the treadmill during this time. Two, two-shank 64-channel silicon probes (NeuroNexus) were advanced through craniotomies to hippocampal areas CA3 and CA1 on each recording day. Recording sites spanned 170 μm. The CA3 probe was advanced vertically through the craniotomy to a depth of ∼2.15mm. The probe was lowered through cortex, through the CA1 pyramidal layer, and the recording location was chosen as the location where large spikes (100+ μV) and SWRs re-emerged. The CA1 probe was advanced at an angle 35° from vertical and 50° from the coronal plane (right hemisphere) or 28° from vertical and 75° from the coronal plane (left hemisphere) until hippocampal pyramidal layer characteristics were observed (large theta waves and sharp wave ripples, 150+ μV spikes on multiple channels). Data were acquired with a sampling rate of 20 kHz using an Intan RHD2000 Evaluation System using a ground pellet placed in the recording chamber on the skull as reference.

### Audio-visual flicker stimulation

After behavioral training, animals were exposed to either 40 Hz or Random audio-visual flicker. Both stimuli consisted of aligned auditory (65 dB) and visual (160-780 lux) pulses of 12.5ms (50% duty cycle of 40 Hz). The auditory pulses were 10 kHz tones. During 40 Hz stimulation, audio-visual pulses turned on and off periodically (12.5ms on-12.5ms off), while during Random stimulation, the inter-pulse interval was chosen from a random distribution with a mean of 12.5ms. Thus, the two experimental groups were exposed to a similar amount of light and sound pulses during the stimulation, but only the 40 Hz group was exposed to a periodic stimulus. The experimenter was blind to the stimulation condition during behavioral training, electrophysiology experiments, flicker exposure, and data preprocessing. Blinding was maintained via procedures that we have described previously (Attokaren et al., 2023). In brief, animals were assigned to the 40 Hz or Random stimulation group by a nonblinded third party. Infrared video was used to monitor animal behavior during the experiment, while blocking the experimenter’s perception of the flicker frequency. The flicker program played a test sequence prior to beginning the 40 Hz or Random stimulation sequence, allowing the experimenter to determine that the light and sound stimuli were working appropriately.

Animals were exposed to 40 Hz or Random stimulation according to their assigned group during recording sessions and during flicker exposure sessions. During electrophysiology recordings, animals were exposed to 1 hour of sensory stimulation while they were free to run or rest on the spherical treadmill (Experimental Days 1, 9, and 10, see Figure 1). The animals were not exposed to the virtual reality environment during flicker. During flicker exposure sessions, animals were placed in a cage similar to their home cage with free access to water and were exposed to audio-visual stimulation for 1 hour (Experimental Days 2-8, see Figure 1). The exposure cage had 3 sides covered with black material, and one side open to face an LED light strip. On Experimental Days 2-8, animals were trained on the VR task to maintain performance. Training occurred at least 1 hour after flicker stimulation.

### Local field potentials (LFP)

Raw traces were downsampled to 2 kHz and bandpass filtered between 1-300 Hz to obtain LFP. Outliers were detected at 15 standard deviations above the mean of the raw signal and were eliminated by interpolation. LFP traces were bandpass filtered using a FIR equiripple filter for delta (1-4 Hz), theta (4-12 Hz), beta (12-30 Hz), and sharp-wave ripples (150-250 Hz). To determine theta periods, the envelope amplitude of the filtered theta signal was divided by the sum of the envelope amplitudes of the delta and beta signals (“theta-deltabeta ratio”) (Csicsvari et al., 1999; Iaccarino et al., 2016). To be classified as a theta period, the theta-deltabeta ratio had to exceed a threshold of 2 standard deviations above the mean for 2 seconds. To determine non-theta periods, the same theta-deltabeta ratio had to be below a threshold of 1.1 for at least 2 seconds. Sharp-wave ripple events were detected when the envelope of the filtered sharp-wave ripple trace was greater than 3 standard deviations above the mean for at least 20 ms (Karlsson & Frank, 2009; Singer et al., 2013; Singer & Frank, 2009). The power ratio of 100-250 Hz to 250-400 Hz was calculated and any events with a power ratio less than 4 were excluded (Ylinen et al., 1995). To exclude any artifacts that may be detected by these parameters, we excluded any events that had an LFP amplitude exceeding ±1500 mV. All detected periods (theta, non-theta, and sharp-wave ripples) were visually inspected to confirm the detection parameters.

Theta and sharp-wave ripple LFP analyses were conducted using the LFP signal from the channel on the electrode with the largest power in the sharp-wave ripple band. This channel is putatively located in the stratum pyramidale and was typically located in the middle of the depth-wise channel span of the probe (Figure S1). We verified that the location of this channel was similar across stimulation conditions by examining the distribution of SWR power across all channels for all recording sessions.

### Spike sorting and cell type classification

Waveform extraction and clustering was conducted using Kilosort2 automated spike sorting software and the Phy2 visualization GUI for manual curation (Pachitariu et al., 2016). After manual curation, units were included for further analysis only if they had less than 0.8% refractory period (1ms) violations and peak signal to noise ratio (SNR) greater than or equal to 1. Isolated single units were then classified into putative pyramidal cells and putative interneurons based on spike width and the first moment of the autocorrelogram (Barthó et al., 2004; Csicsvari et al., 1998, 1999; Niell & Stryker, 2008; Prince et al., 2021; Senzai et al., 2019). Units with a spike width greater than 0.5 ms and a first moment of the autocorrelogram less than 4.5 ms were classified as putative pyramidal cells. Units that had a spike width less than 0.5 ms and a first moment of the autocorrelogram greater than 4.5 ms were classified as putative interneurons. This is similar to previously reported values for putative classification (Csicsvari et al., 1998, 1999; Prince et al., 2021; Senzai et al., 2019).

### Spike-phase coupling

Pairwise phase consistency (PPC) was used to quantify spike-field phase synchronization between LFP and spikes. PPC is unbiased by the number of trials and is less affected by the number of recorded spikes than conventional phase-locking analyses. In short, PPC for a given frequency band *f* is calculated with the following equation (for details refer to: (Vinck et al., 2012a)).

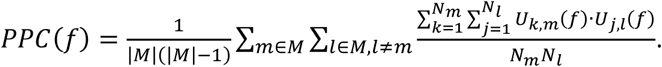

in which |*M*| is the number of trials in total, *U*_*k,m*_ is the instantaneous phase of filtered LFP at frequency *f* when the *k*th spike occurs during trial *m*. *N*_*m*_ and *N*_*l*_ are the number of spikes in trial *m* and *l* respectively. The instantaneous phase of the filtered LFP was calculated using the Hilbert transform. To exclude the impact of flicker-entrained oscillations during flicker, a notching filter of 40 Hz with 1Hz bandwidth was applied to LFPs before above PPC calculation. Neurons included in the spike field analysis had to satisfy the following criteria: neurons fired in at least five trials with a total of more than one hundred spikes to ensure that enough spikes were available for analysis. During VR behavior periods, trials were defined from the end of one reward zone through the next reward zone, while trials during flicker stimulation were defined as 1-minute-long periods during recording.

### CA3-CA1 weighted phase lag index

Weighted phase lag index (WPLI) was calculated to measure CA3-CA1 phase-synchronization. WPLI is impacted less by spurious synchrony caused by common reference, volume-conduction, common noise, and sample-size bias than most other existing methods (Vinck et al., 2012). To calculate WPLI, first the cross-spectrum between CA3 and CA1 LFPs is computed as *X*_*CA*3−*CA*1_, using Welch’s method with a non-overlapped Hamming window of 1024 points and NFFT = 1024. Then we calculated WPLI as following:

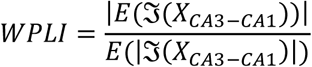

where ξ(*X*_*CA*3−*CA*1_) is the imaginary component of *X*_*CA*3−*CA*1_, and *E*(·) represents the mean. Calculation of WPLI was performed using modified code from the FieldTrip toolbox (https://github.com/fieldtrip/fieldtrip/tree/master) in MATLAB (Oostenveld et al., 2011).

### Place cell identification

Occupancy-normalized firing rate maps were calculated for all putative pyramidal cells. Firing rate was calculated as a function of position over the full track span (360 degrees) using 2-degree spatial bins to calculate spike count and time spent in each spatial bin. Only times and spikes where the animal was moving (speed > 1 deg/s) were included. Spiking and occupancy maps were smoothed using a Gaussian window with a standard deviation of 2 spatial bins. Occupancy-normalized firing rate maps were computed by dividing the smoothed spiking by the smoothed occupancy in each position bin. Spatial information was calculated for each unit using the following equation:

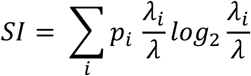

where SI is the spatial information content of the unit in (bits/spike), *p*_*i*_ is the probability of the animal being in the *i*th spatial bin, λ_*i*_ is the mean firing rate of the unit in the *i*th spatial bin, and λ is the mean firing rate of the unit across all spatial bins (Langston et al., 2010; Skaggs et al., 1996). Spatial information for each unit was compared to the spatial information calculated for shuffled spike train data. To compute shuffled spike trains, each spike was permuted by a random time from 20s to the length of the recording session. Any spike times greater than the length of the recording were wrapped around to the beginning of the recording session. Shuffled firing rate maps and spatial information values were calculated from these shuffled spike trains. The shuffling process was repeated 500 times for each pyramidal cell. Units with real spatial information values greater than the 95^th^ percentile of the shuffled spatial information values were included for further analysis. Putative pyramidal cells were classified as place cells if they met the spatial information criteria detailed above, had a minimum mean firing rate of 0.2 Hz, had a maximum mean firing rate of 10 Hz, and had a minimum peak firing rate of 1 Hz. The same criteria were applied to putative pyramidal cells from CA1 and CA3. Because behavior was similar across both halves of the track, we performed many analyses as a function of distance to the rewarded zones. Once place cells were identified over the full track, we recalculated the rate maps for these cells as a function of distance to the reward zone using 3-degree spatial bins. For place field visualization, occupancy-normalized firing rate maps are shown with as the occupancy-normalized firing rate normalized by the mean firing rate.

### Place cell properties

Place cell properties were calculated from the distance to reward occupancy-normalized firing rate maps computed from place cells classified as detailed above. Sparsity of each rate map was calculated using the following equation:

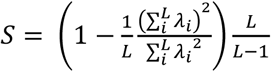

Where S is the sparsity of the place cell rate map (S = 0 indicates a uniform rate map, S = 1 indicates a unidirectional rate map), L is the total number of spatial bins (L = 60 for distance to reward rate maps), and λ_*i*_ is the mean firing rate of the unit in the *i*th spatial bin (Ravassard et al., 2013). The maximum firing rate of the unit was the maximum of the unit’s occupancy-normalized firing rate map. The peak firing position was the position within the track where the maximum of the occupancy-normalized firing was located. The mean firing rate of the place cell was the average of the occupancy-normalized firing rate map.

### Bayesian decoding of current position

Estimated position was decoded from place cell spiking activity using a simple Bayesian decoder (Brown et al., 1998; Karlsson & Frank, 2009). Correct trials (1 trial is a single pass through a reward zone, 180°, ½ lap around the track) for each recording session were randomly divided into training and testing datasets using 5-fold cross validation. The training dataset was used to calculate occupancy-normalized firing rate maps with 3-degree spatial bins for all place cells. To compute position estimates, spikes from testing trials were binned into temporal blocks of 200 ms. For each temporal block with at least 2 spikes, the spatial probability estimate was calculated using the following equation:

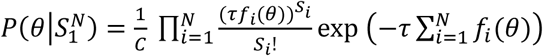

where *N* is the number of included cells, τ is the time window to be decoded, θ is position, *S*_*i*_ is the spiking activity of cell *i* in the time window, *f*_*i*_(θ) is the occupancy-normalized firing rate for cell *i* over position θ, and *C* is a constant used to normalize *P*(θ|*S*^*N*^) to sum to 1. Position estimates for each temporal block were binned and averaged by the animal’s actual position (the average track position during the temporal block). Decoding error was calculated as the difference between the spatial location with the highest estimated probability and the animal’s actual position. The decoding result was cross-validated (5-fold cross validation) and the output was averaged. To ensure analyses were only performed on sessions with acceptable decoder performance, sessions with an averaged normalized probability of decoding actual position less than 1.2 were excluded from current position and sharp-wave ripple (see *Sharp-wave ripple decoding*) decoding analyses.

### Theta sequences

Correct trials (trials where animals licked for a reward) were used to calculate theta sequences. During these trials, individual theta cycles were identified during periods of running. The phase of theta was calculated using the Hilbert transform. A 400 ms window was centered around each theta trough, and place cell spiking during this window was divided into 20 ms temporal bins (Farooq & Dragoi, 2019). A Bayesian decoder (see *Bayesian decoding of current position*) was trained on the occupancy-normalized firing rate maps of place cells, and the decoder was used to calculate spatial probability estimates for each temporal bin during the theta cycle window. Only bins with 2 or more spikes were decoded. The resulting probability estimate was centered on the animal’s actual position in the track (position at the theta trough). Decoded windows from the entire recording session were averaged together and the quadrant ratio was computed to determine the strength of the sequence:

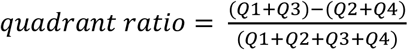

where Q1 represents the future quadrant (0-80 ms, 0-45 degrees) and Q3 represents the past quadrant (−80-0 ms, −45-0 degrees). To determine the significance of the sequence, the time bins of the decoded theta cycles were shuffled, and the quadrant ratio was recalculated 500 times. The recording session was considered to have significant theta sequences if the quadrant ratio of the session average was greater than the 95^th^ percentile of the shuffled quadrant ratios. Only recording sessions with significant theta sequences were included for further analysis.

### Prospective coding during theta

To compute the ratio of prospective to retrospective information represented within the theta sequence, we defined a prospective coding ratio based on the future and past quadrants of the theta sequence:

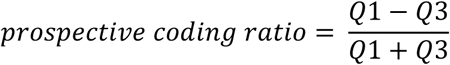

The prospective coding ratio will thus have a value of 1 if the theta sequence represents only prospective information and a value of −1 if only retrospective information is represented. We computed the prospective coding ratio on average theta sequences across the track. To do this, we binned the adjusted decoded theta windows by the animal’s position during the window, averaged the windows in each bin, and used these averages to compute the prospective coding ratio over position.

To analyze prospective coding during trials with different running speeds, the average velocity outside of the last half of the anticipatory zone and the reward zone was computed for each trial. Median trial speed was computed from all correct trials from both stimulation groups then that median speed was used to split the trials into the top half of trial speeds (fast trials) and the bottom half of trial speeds (slow trials).

### Unoccupied reward zone decoding during theta

Decoding of the unoccupied reward or control zone was determined by performing Bayesian decoding of the full 360° track during theta cycles (see *Theta Sequences*). Place cell firing rate maps used in the decoder were computed over the full 360° track. Only recording sessions with significant theta sequences as determined by the previous 180° sequence analysis were included. Decoded theta cycles were averaged per trial as a function of position in the reward zone or control zone. These averaged theta cycles were determined to contain representation of the unoccupied zone if the decoded probability 162-198° away from the animal’s current position was greater than or equal to 2 standard deviations above the mean of the entire cycle. When the animal was in the reward zone, this approach identified decoding of the other, unoccupied reward zone because the track contains two reward zones 180 degrees apart.

When the animal was in the control zone, this approach identified decoding of a control zone away from the reward zone.

### Sharp-wave ripple decoding

Position content during sharp-wave ripples was estimated using a simple Bayesian decoder (see *Bayesian decoding of current position*) trained on occupancy-normalized place cell firing rate maps. First, we decoded the place cell spiking activity during each SWR using multiple time bins. Place cell spiking activity in a 250ms window around each SWR mid-point was divided into 25ms temporal bins. A spatial probability estimate was computed for each temporal bin with more than 1 spike. To quantify the amount of prospective or retrospective information in each SWR, we calculated the spatial probability estimates relative to the animal’s actual position. Then, we computed the sum of the estimated spatial probabilities in prospective positions (+9 to +63 degrees in front of the animal) and the sum of the estimated spatial probabilities in retrospective positions (−9 to −63 degrees behind the animal). Local positions (+/- 9 degrees, equivalent to 1 zone in the track) were excluded to exclude coding of very local positions. The edges of the track (−63 to −90 and +63 to +90) were excluded from this calculation due to the circular nature of the track. The prospective coding ratio for SWRs was calculated as the difference between the sum of the estimated spatial probabilities in the prospective window and the sum of the estimated spatial probabilities in the retrospective window, normalized by the sum of the estimate spatial probabilities in both windows. Thus, a prospective coding ratio value of 1 indicates that the SWR only contains information about locations in front of the animal’s real position, and a prospective coding ratio value of −1 indicates that the SWR only contains information about locations behind the animal’s real position.

### Unoccupied reward zone decoding during sharp-wave ripples

Decoding of the unoccupied reward zone during SWRs was determined by performing Bayesian decoding of the full 360° track during SWR events that occurred in the reward zone (see *Sharp-wave Ripple Decoding*). Place cell firing rate maps used in the decoder were computed over the full 360° track. Positions were estimated in 25 ms temporal bins around the SWR mid-point. These position estimates were used to determine if the SWR contained representation of the unoccupied reward zone. If the decoded probability 180° away from the animal’s current position was greater than or equal to 2 standard deviations above the mean of the entire decoded SWR event, the SWR was considered to have decoding of the unoccupied reward zone.

### Sharp-wave ripple activation and coactivation

Place cell activation probabilities were calculated by determining the number of SWR events in which the cell fired at least one spike, divided by the total number of SWR events on that recording day. Place cell pair coactivation was determined as the number of SWRs in which a pair of place cells both fired, divided by the total number of SWRs on that recording day.

### Statistical analysis

Animals from the 40 Hz and Random stimulation groups each contributed 1-2 recording sessions of data, and each recording session contributed a distinct number of single units, trials, and sharp-wave ripple events. To account for animal-specific effects, we fitted a linear-mixed effect model (LME) using the *lme4* package in R (Bates et al., 2015; R Core Team, 2022). The LME approach assesses the significance of differences between groups while controlling for repeated measures from the same animals and sessions. Using linear-mixed effects models to account for repeated measures within an animal is advantageous over computing a single value, like an average, per animal as there is often high variation in cells or patterns of activity recorded within an animal and thus an average may not accurately represent the distribution of the data. The model included the stimulation group as the fixed effect and animal as the random effect. A F test using the Kenward-Roger method to calculate the denominator degrees of freedom was performed to determine significance. Throughout, error bars are shown as mean±SEM. Percent changes and fold changes between measures from 40 Hz and Random groups were calculated as the difference between groups divided by the absolute value of the Random group and multiplied by 100 for percentages. Statistical details and further quantification of all results is available in Supplementary Tables 2-4. Significance values are reported as follows: +P<0.1,*P<0.05, **P<0.01, ***P<0.001, ****P<0.0001.

## Data Availability

Analysis code is available at https://github.com/abigailpaulson/flicker-neuralcodes/

## Acknowledgements

We thank Laura Colgin and the Singer lab for valuable comments on the manuscript. A.C.S. was supported by the Packard Foundation, the National Institutes of Health (NIH)-National Institute of Neurological Disorders and Stroke Grant R01 NS109226 and 2RF1NS109226, the NIH National Institute of Aging Grant RF1AG078736, McCamish Foundation, Friends and Alumni of Georgia Tech, and the Lane Family. A.L.P. was supported by the NIH National Research Service Award Grant 5F31AG066410, the NIH Grant T32 NS007480-18, Fulton County Elder Health Science Fellowship, the Wright Family, and J. Norman and Rosalyn Wells Fellowship. A.M.P. was supported by Veterans Affairs (VA) grant CDA2 E4544-W.

## Author contributions

Conceptualization, A.L.P. and A.C.S.; Methodology, A.L.P and A.C.S.; Investigation, A.L.P., A.M.P., and L.Z.; Formal Analysis, A.L.P. and L.Z.; Data Curation, A.L.P. and A.M.P.; Writing – Original Draft, A.L.P. and A.C.S.; Writing – Review and Editing, A.L.P, L.Z., A.M.P., and A.C.S.; Visualization, A.L.P., L.Z., and A.C.S.; Funding Acquisition, A.C.S.; Supervision, A.C.S.

## Additional Files

### Additional File 1

**.mov**

**A headfixed mouse performs the virtual reality spatial navigation task.**

The mouse runs on the spherical treadmill, which controls movement through the virtual reality track. The mouse must lick in specific zones of the track to receive a reward of sweetened condensed milk.

