## Supplementary Information for "40 Hz sensory stimulation enhances CA3-CA1 coordination and prospective coding during navigation in a mouse model of Alzheimer’s disease"

Supplement Includes:

- Supplementary Figures 1-10
- Supplementary Tables 1-6

**Supplementary Figures**

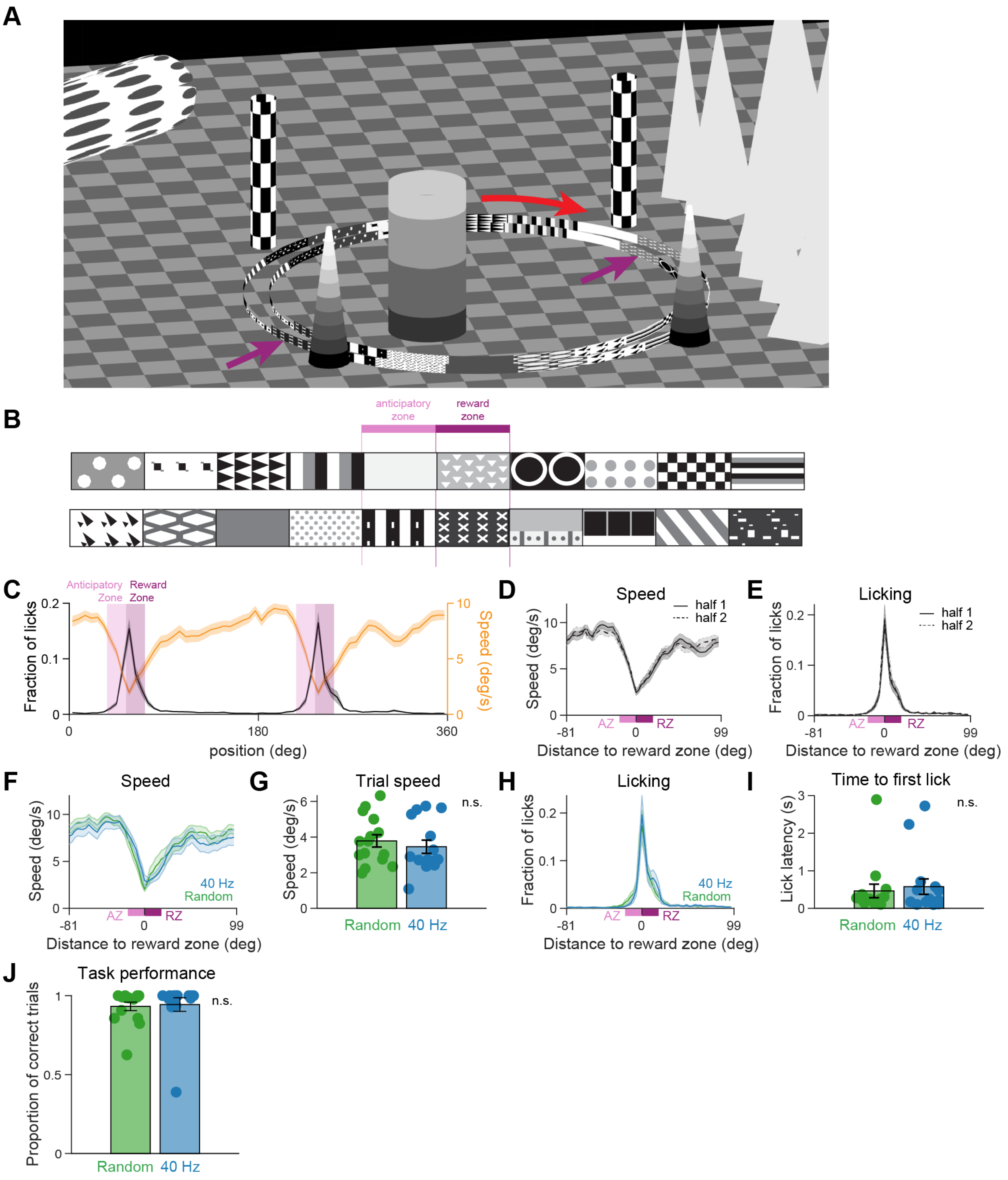


**Supplementary Figure 1. 5XFAD mice perform similarly in a goal-directed spatial navigation task after 8 days of 40 Hz or Random stimulation.**

**A.** Overhead view of the virtual reality environment. The mouse runs through the annular track in the direction indicated by the red arrow. The reward zones are indicated with purple arrows

(arrows are not part of the environment and are not shown to the animal). Distal cues (pillars, floating objects) are visible above the patterned walls as the mouse runs through the track.

**B.** Detailed schematic of the patterned cues in the track centered around the anticipatory and reward zones. Each patterned cue covers 18 degrees of the 360 track.

**C.** Average speed (orange, right y-axis) and licking (black, left y-axis) as a function of position in the annular environment. 360 degrees indicates a full traversal of the track. Reward zones, located 180 degrees apart, are indicated by dark pink shaded areas with anticipatory zones just before reward zones indicated by light pink shaded areas. Mean ± SEM across recording days. Random, n = 15 days in 8 mice; 40 Hz, n = 15 days in 8 mice.

**D.** Speed as a function of position is similar between the two halves of the track (solid line, first half and dashed line, second half). Animals display similar slowing behavior in the anticipatory zone (AZ, light pink) as they approach both reward zones (RZ, dark pink). Mean ± SEM across recording days. Random, n = 15 days in 8 mice; 40 Hz, n = 15 days in 8 mice.

**E.** As animals approach the rewarded areas of the track, licking increases in anticipation of the reward. When the animals are in the reward zone (dark pink), increased licking indicates reward consumption. Anticipatory licking behavior occurs before both reward zones in the anticipatory zone (light pink). Mean ± SEM across recording days. Random, n = 15 days in 8 mice; 40 Hz, n = 15 days in 8 mice.

**F.** Animals exposed to 40 Hz (blue) and Random (green) flicker show anticipatory slowing behavior as they approach the reward zone. Mean ± SEM across recording days. Random, n = 15 days in 8 mice; 40 Hz, n = 15 days in 8 mice.

**G.** Average trial speed per day for animals exposed to 40 Hz (blue) and Random (green) flicker. Mean ± SEM across recording days. 40 Hz, n = 15 days in 8 mice; Random, n = 15 days in 8 mice, P=0.51, n.s., linear mixed-effects model (LME). See Supplementary Table 3 for statistical details.

**H.** As in **D** for licking behavior. Animals exposed to 40 Hz and Random flicker show anticipatory licking behavior as they approach the reward zone. Random, n = 15 days in 8 mice; 40 Hz, n = 15 days in 8 mice. Mean ± SEM across recording days.

**I.** Average time to first lick in the rewarded zones of the track. Most animals lick quickly (<1 s) following entry into the reward zone. Reward is not delivered until animals lick inside the rewarded zone. Mean ± SEM across recording days. 40 Hz, n = 15 days in 8 mice; Random, n = 15 days in 8 mice, P=0.74, n.s., LME.

**J.** Task performance was not significantly different between animals exposed to 40 Hz or Random flicker. Mean ± SEM across recording days. 40 Hz, n = 15 days in 8 mice; Random, n = 15 days in 8 mice, P=0.85, n.s., LME.

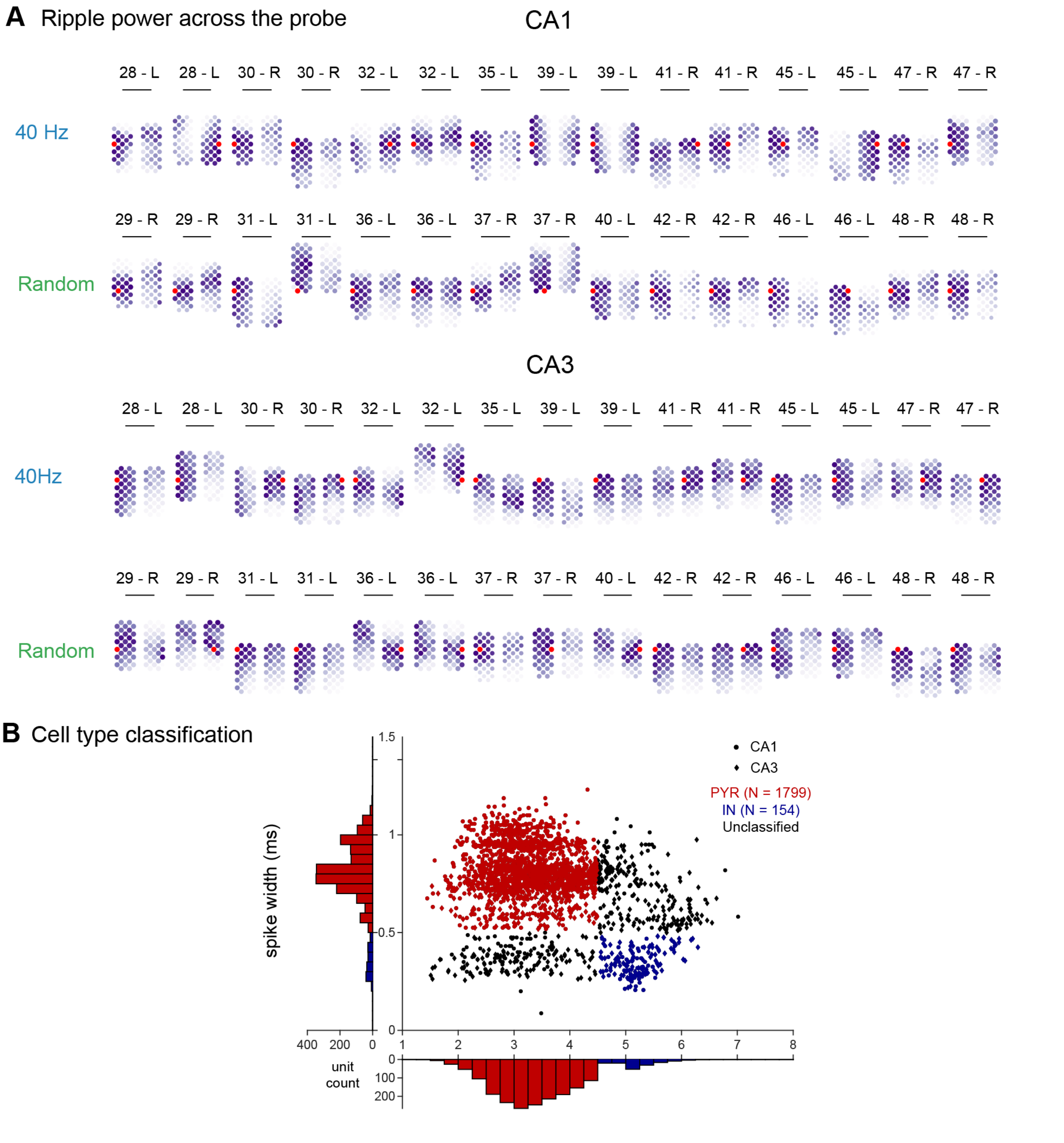


**Supplementary Figure 2. Recording locations and cell type classification.**

**A.** Recording location for all recording sessions in CA1 (top two rows) and CA3 (bottom two rows) is shown by quantifying the ripple power (150-250 Hz) across the probe. The electrode has two shanks, each with 32 channels. Each circle is a recording channel. The color of the channel corresponds to the ripple power recorded on the channel; darker purple colors indicate higher ripple power with the red channel indicating the largest ripple power channel on each recorded day (used for ripple analyses, see Methods). The ripple power is expected to be highest in the center of the pyramidal cell layer. Probe diagrams are aligned to the channel with the highest ripple power. The animal ID and recording hemisphere (left (L) or right (R) ) are noted above each probe illustration. Two recording sessions were performed from each hemisphere.

**B.** Recorded single units were classified into putative pyramidal cells (red) and putative interneurons (blue) based on spike width and the mean of the autocorrelogram. The same criteria were applied to CA1 units (circles) and CA3 units (diamonds). Unclassified cells are marked in black and were excluded from further analyses. CA1, n = 1011 pyramidal cells, n = 54 interneurons; CA3, n = 788 pyramidal cells, n = 100 interneurons.

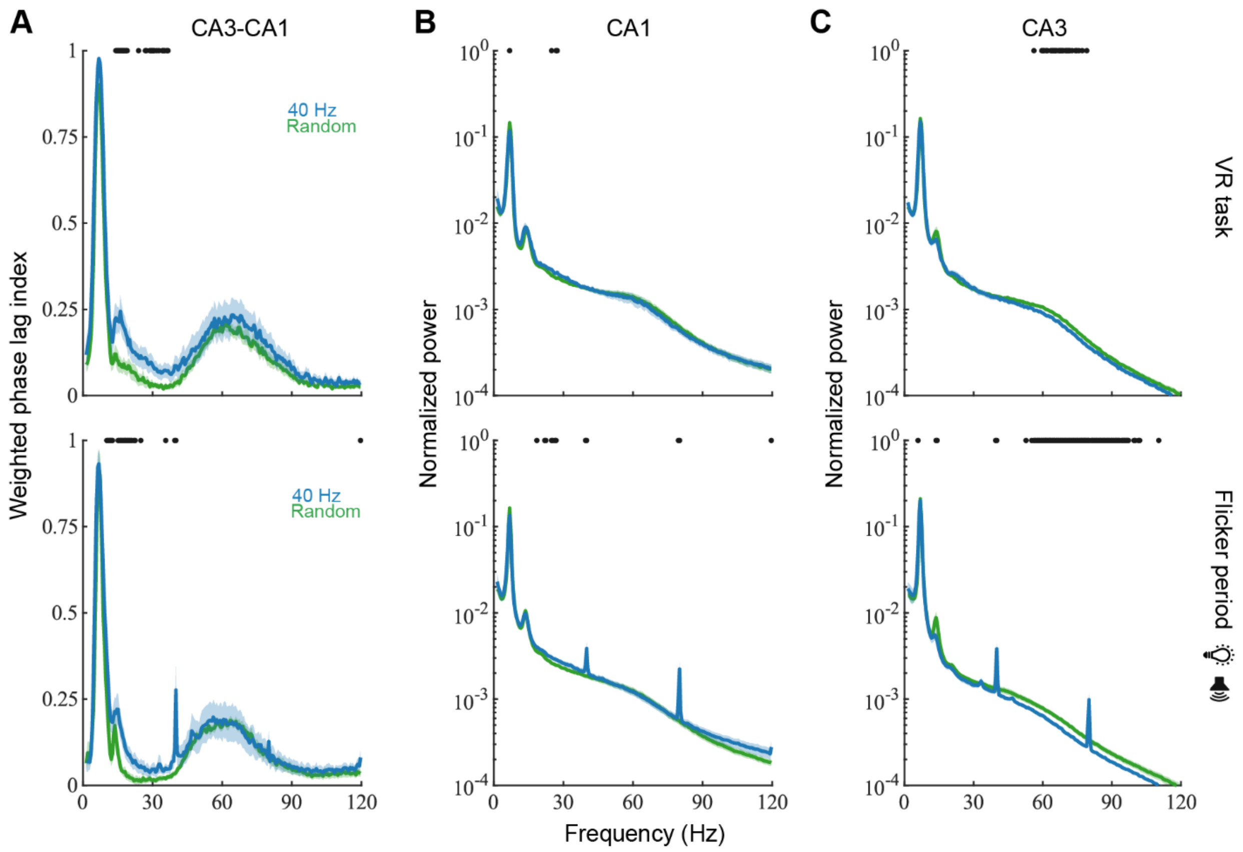


**Supplementary Figure 3. CA3-CA1 LFP phase synchrony is enhanced in the slow gamma band following 40 Hz flicker.**

- **A.** Weighted phase lag index between LFPs recorded in CA3 and CA1 during VR navigation periods (top) and flicker stimulation periods (bottom) after exposure to 40 Hz (blue) or Random (green) flicker. Weighted phase lag index measures functional connectivity with minimal volume conduction effects. Functional connectivity between CA3 and CA1 was higher after 40 Hz than Random flicker in slow gamma bands. Dots above the graph indicate frequencies with significant differences between 40 Hz and Random flicker exposure groups. $p < 0.05$, unpaired t-test without correction, 40 Hz, n = 15 days in 8 mice; Random, n = 15 days in 8 mice. See Supplementary Table 4 for statistical details. Mean ± SEM across recording days.
- **B.** Normalized power spectral density of LFPs recorded in CA1 during VR navigation task periods (top) and flicker stimulation periods (bottom) after exposure to 40 Hz (blue) or Random (green) flicker. Dots indicate frequency points with significant differences between 40 Hz and Random ($p < 0.05$, unpaired t-test without correction, 40 Hz, n = 15 days in 8 mice; Random, n = 15 days in 8 mice). Mean ± SEM across recording days.
- **C.** Same as **B** for LFPs in CA3.

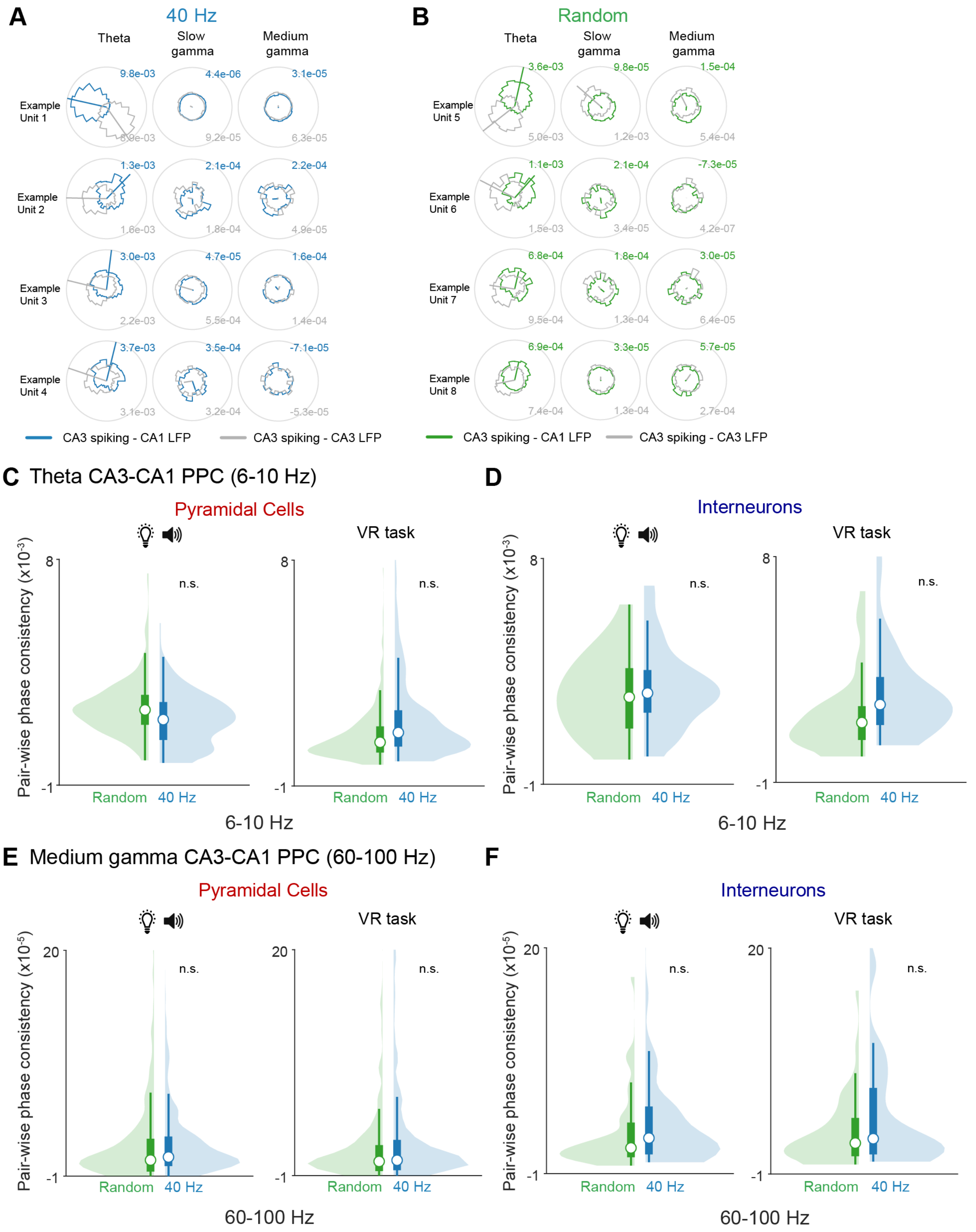


**Supplementary Figure 4. Theta and medium gamma PPC are similar following 40 Hz or Random flicker.**

**A.** Local (grey) and cross-region (blue) spike-phase coupling for example single units in CA3 and LFPs in CA1 during periods of theta (left), slow gamma (center), and medium gamma (right) after exposure to 40 Hz flicker. Each row is an example single unit.

**B.** Same as **A** for example CA3 single units from animals exposed to Random stimulation with local coupling in grey and cross-region coupling in green.

**C.** Theta (6-10 Hz) spike-field pairwise phase consistency (PPC) for putative CA3 pyramidal cells and CA1 LFPs during flicker stimulation periods (left) and during VR behavior periods (right) following 8 days of exposure to 40 Hz (blue) or Random (green) flicker stimulation. *Left*, flicker stimulation periods, Random, n = 320 pyramidal cells from 8 mice; 40 Hz n = 387 pyramidal cells from 8 mice; P = 0.19, n.s., linear mixed-effects model (LME). *Right*, VR task periods, Random, n = 347 pyramidal cells from 8 mice; 40 Hz, n = 335 pyramidal cells from 8 mice; P = 0.29, n.s., LME. See Supplementary Table 3 for statistical details. Violin plots throughout include a histogram (shaded area) and box plot indicating median (white circle), first and third quartiles (dark box) and whiskers (thin lines).

**D.** As in **B** for putative CA3 interneurons. *Left*, flicker stimulation periods, Random, n = 46 interneurons from 8 mice; 40 Hz, n = 50 interneurons from 8 mice; P = 0.36, n.s., LME. *Right,* VR task periods, Random, n = 49 interneurons from 8 mice; 40 Hz, n = 50 interneurons from 8 mice; P = 0.12, n.s., LME.

**E.** Medium gamma (60-100 Hz) spike-field pairwise phase consistency (PPC) for putative CA3 pyramidal cells and CA1 LFPs during flicker stimulation periods (left) and during VR behavior periods (right) following 8 days of exposure to 40 Hz (blue) or Random (green) flicker stimulation. *Left*, flicker stimulation periods, Random, n = 320 pyramidal cells from 8 mice; 40 Hz n = 387 pyramidal cells from 8 mice; P = 0.90, n.s., linear mixed-effects model (LME). *Right*, VR task periods, Random, n = 347 pyramidal cells from 8 mice; 40 Hz, n = 335 pyramidal cells from 8 mice; P = 0.32, n.s., LME.

**F.** As in **E** for putative CA3 interneurons. *Left*, flicker stimulation periods, Random, n = 46 interneurons from 8 mice; 40 Hz, n = 50 interneurons from 8 mice; P = 0.21, n.s., LME. *Right,* VR task periods, Random, n = 49 interneurons from 8 mice; 40 Hz, n = 45 interneurons from 8 mice; P = 0.11, n.s., LME.

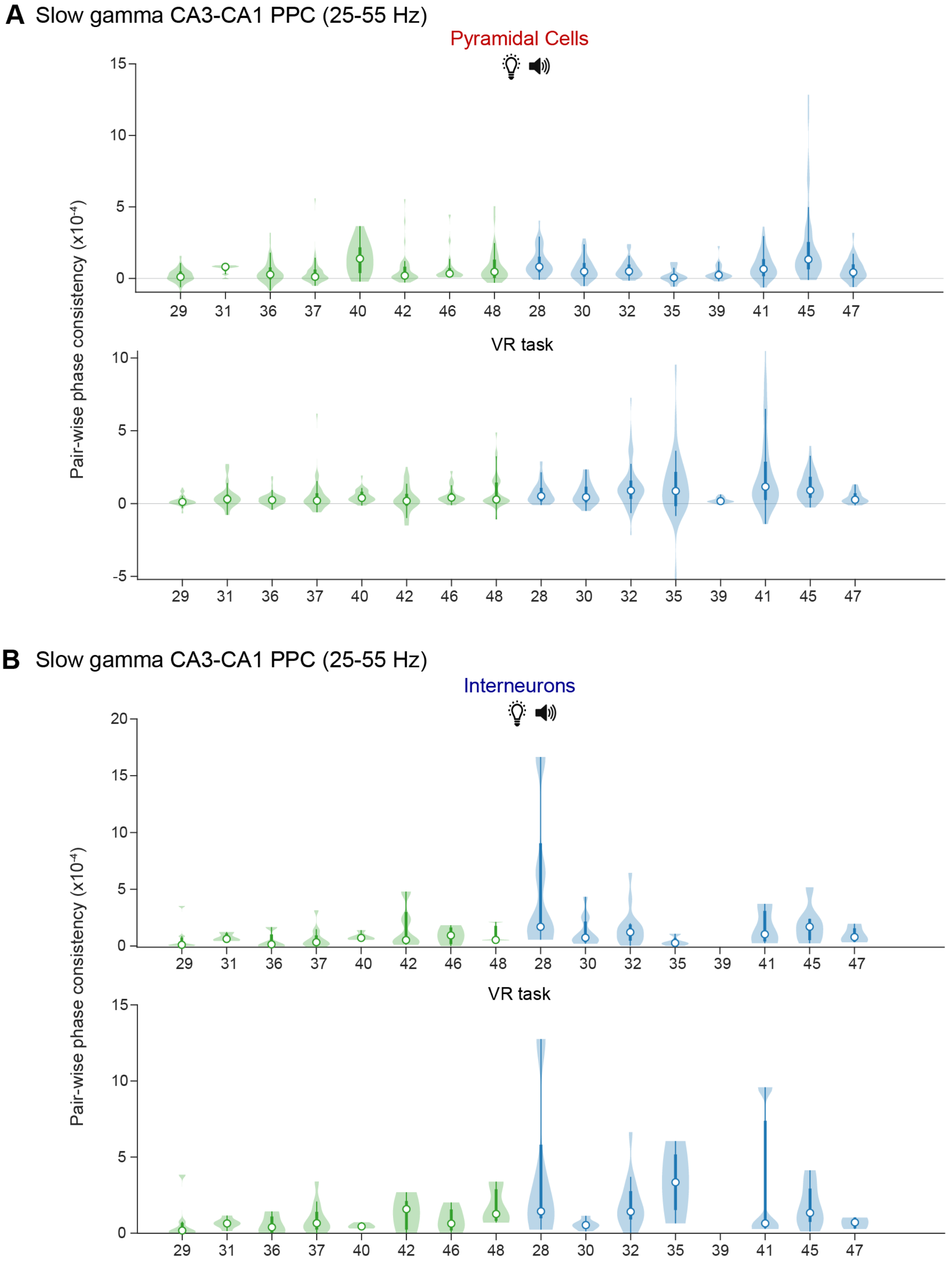


**Supplementary Figure 5. Slow gamma CA3-CA1 pairwise phase consistency distributions per animal.**

**A.** Slow gamma (25-55 Hz) pairwise phase consistency between spiking from putative CA3 pyramidal cells and CA1 LFP during flicker exposure (top) and VR behavior (bottom) periods per animal. Each violin plot contains data from one animal. The animal ID is noted on the x-axis.

**B.** Same as in **A** but for putative CA3 interneurons.

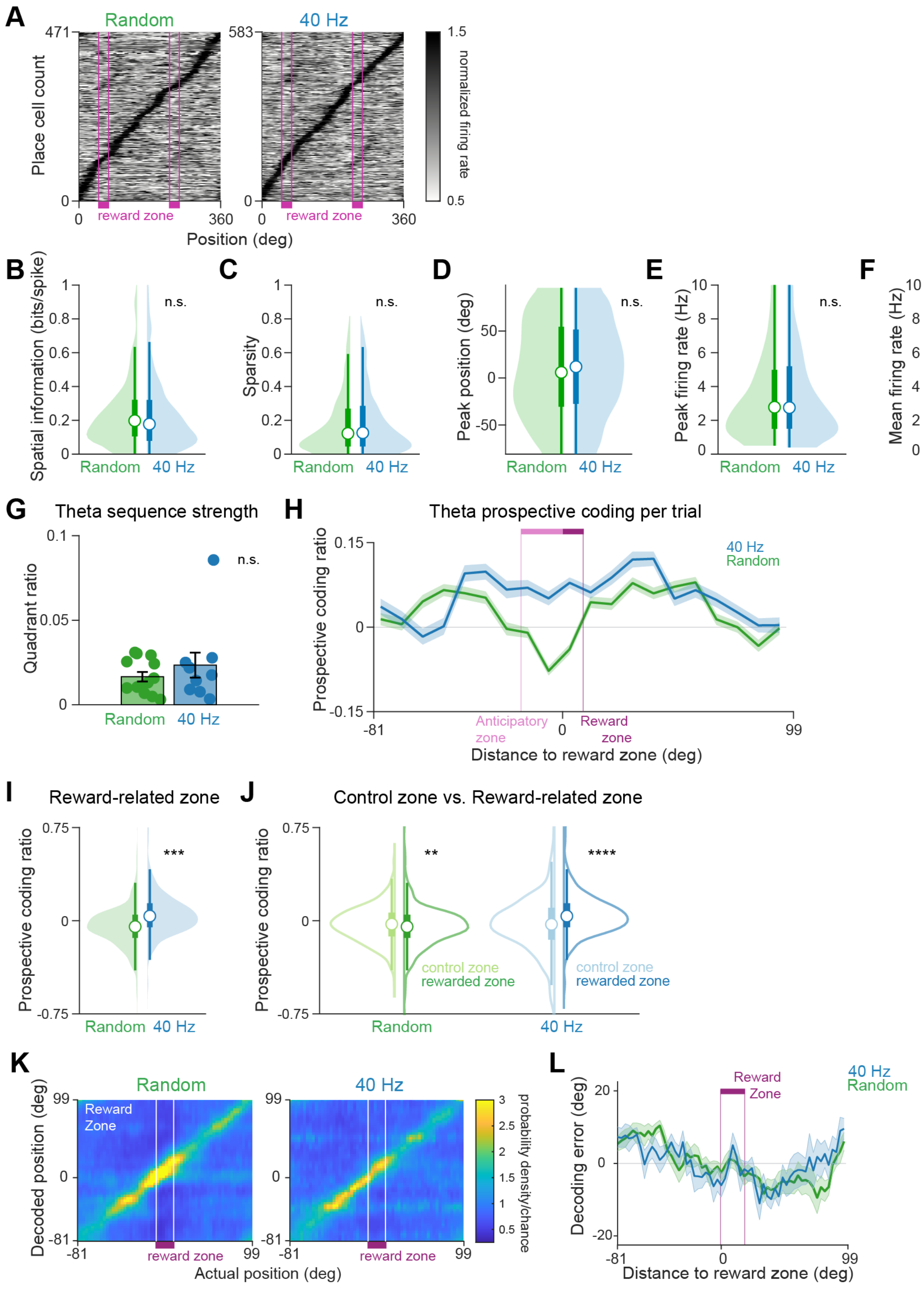


**Supplementary Figure 6. Place cell properties and decoding performance are similar following 40 Hz and Random flicker.**

- **A.** Normalized firing rate maps for place cells recorded from animals exposed to Random (left) or 40 Hz (right) stimulation with each row indicating a single place cell. Cells are ordered by peak firing position in the track. Reward zones are indicated by pink lines and boxes along the x-axis. Random, n = 471 place cells from 8 mice; 40 Hz, n = 583 place cells from 8 mice.

**B.** Distribution of spatial information of place cells shown in **A** following 8 days of 40 Hz (blue) or Random (green) flicker exposure. Spatial information was computed from rate maps that were calculated as a function of distance to the reward zone in the track, as pictured in **Figure 3G**. Random, n = 471 place cells from 8 mice; 40 Hz, n = 583 place cells from 8 mice, P = 0.93, n.s.; linear mixed-effects model (LME). See Supplementary Table 3 for statistical details.

**C.** As in **B** for place cell sparsity, which ranges from 0 (the cell fires equally in all spatial bins) to 1 (the cell only fires in one spatial bin) to assess the relative proportion of the track in which the cell fired. Random, n = 471 place cells from 8 mice; 40 Hz, n = 583 place cells from 8 mice; P = 0.69, n.s., LME.

**D.** As in **B** for peak firing position of place cells as a function of distance to the reward zone. Random, n = 471 place cells from 8 mice; 40 Hz, n = 583 place cells from 8 mice; P = 0.68, n.s., LME.

**E.** As in **B** for maximum firing rate of place cells. Random, n = 471 place cells from 8 mice; 40 Hz, n = 583 place cells from 8 mice; P = 0.86, n.s., LME.

**F.** As in **B** for average firing rate of place cells. Random, n = 471 place cells from 8 mice; 40 Hz, n = 583 place cells from 8 mice; P = 0.85, n.s., LME.

**G.** Average theta sequence strength per day following 8 days of 40 Hz (blue) or Random (green) flicker exposure. Mean $\pm$ SEM across recording days. Random, n = 13 days from 8 mice; 40 Hz, n = 10 days from 6 mice; P = 0.37, n.s., LME.

**H.** Prospective coding ratio over position, calculated per trial. 8 days of 40 Hz flicker (blue) exposure lead to more prospective coding (positive values) in and around the anticipatory zone (light pink) and reward zone (dark pink) than Random flicker (green). Mean $\pm$ SEM across trials. Random, n = 934 trials from 8 mice; 40 Hz, n = 807 trials from 6 mice.

**I.** Per trial prospective coding ratio values in the anticipatory and reward zones were higher following 8 days of 40 Hz (blue) than Random (green) flicker exposure. Random, n = 934 trials from 8 mice; 40 Hz, n = 807 trials from 6 mice; P = 0.0007***, LME.

**J.** Prospective coding ratio was significantly higher after 40 Hz flicker in the reward-related areas (dark colors) of the track than a control location (light colors), where the animal did not receive reward. Random, control zone, n = 931 trials, reward-related zone, n = 934 trials, P = 0.009**, LME. 40 Hz, control zone, n = 807 trials, reward-related zone n = 799 trials, P = 1.4e-13****, LME.

**K.** Actual position of the animal versus Bayesian decoding of current spatial position based on neural activity, averaged across days shows strong decoding of current position following 8 days of 40 Hz (blue) or Random (green) flicker exposure. This validates our decoding approach. Warmer colors along the x=y diagonal indicate accurate neural decoding of the animal's current actual position. The reward zone is indicated by the white lines. Random, n = 13 days from 7 mice; 40 Hz, n = 13 days from 7 mice.

**L.** Decoding error between true and decoded position across the track, averaged across days. Values near zero indicate accurate neural decoding of the animal's current actual position. Decoding accuracy was high, especially approaching and in the reward zone (pink). Mean $\pm$ SEM across recording days. Random, n = 13 days from 7 mice; 40 Hz, n = 13 days from 7 mice.

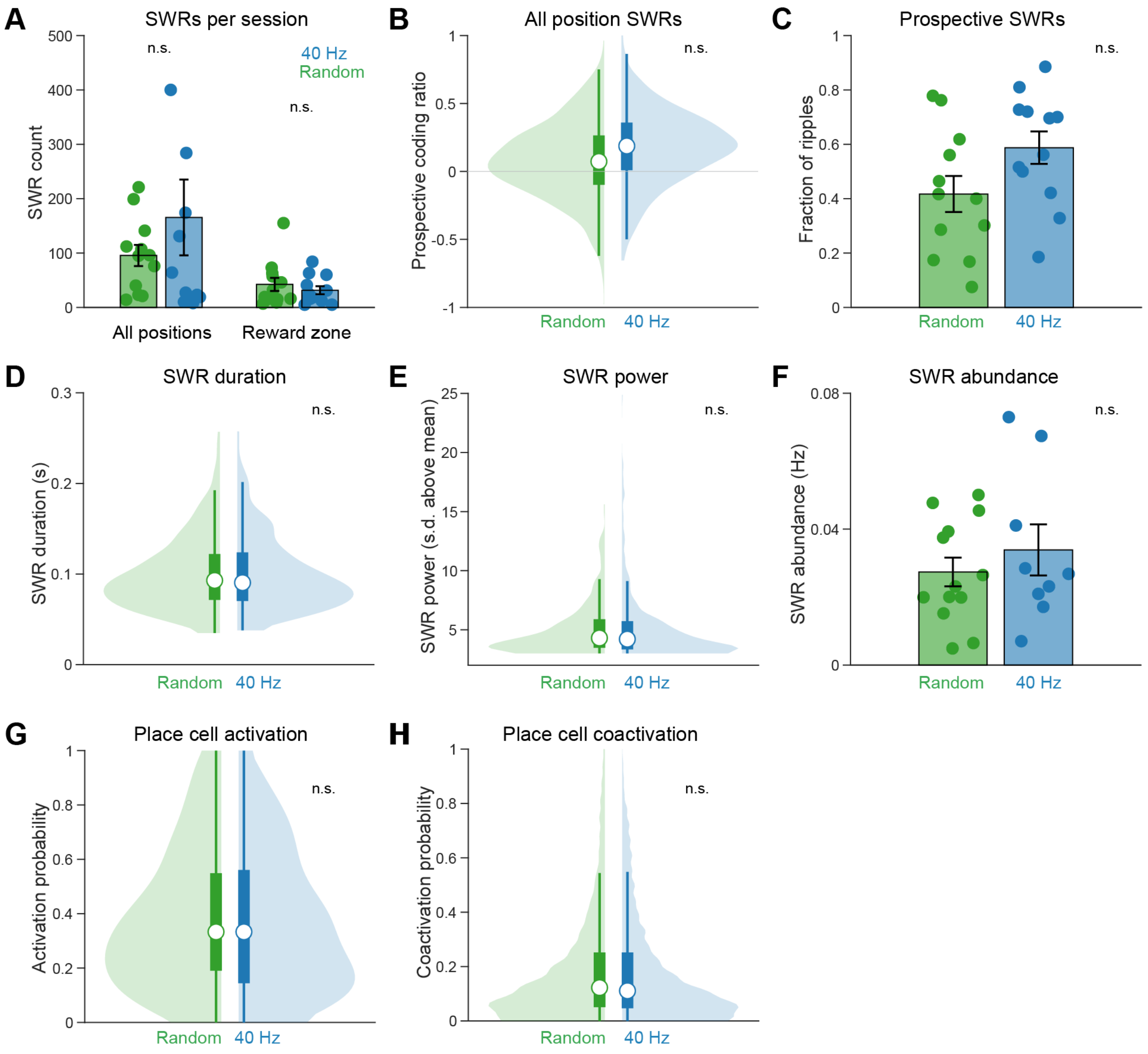


**Supplementary Figure 7. Sharp-wave ripple properties are similar following 40 Hz and Random flicker.**

- **A.** Number of SWRs detected per session for SWRs occurring at all track positions (left) and SWRs occurring inside of the reward zone (right) did not differ significantly following 8 days of 40 Hz (blue) or Random (green) flicker. Only sessions with more than 5 SWRs and with significant decoding (see Methods) were included. Mean ± SEM across recording days. *Left,* SWRs occurring in any position in the track, Random, n = 12 days from 7 mice; 40 Hz, n = 12 days from 7 mice; P = 0.46, n.s., linear mixed-effects mode (LME). *Right*, SWRs occurring in the reward zone of the track, Random, n = 12 days from 7 mice; 40 Hz, n = 12 days from 7 mice; n.s., LME. See Supplementary Table 3 for statistical details.
- **B.** Distribution of prospective coding ratio values for all SWRs occurring at any position in the track did not differ significantly following 8 days of 40 Hz (blue) or Random (green) flicker, unlike SWRs in the reward zone (**Fig. 4C**). Random, n = 1145 SWRs from 7 mice; 40 Hz, n = 1989 SWRs from 7 mice; P = 0.21, n.s., LME.
- **C.** Related to **B,** fraction of SWRs occurring in any location with prospective coding ratios greater than 0.1 did not differ significantly following 8 days of 40 Hz (blue) or Random (green) flicker,

unlike SWRs in the reward zone (**Fig. 4E**). Mean ± SEM across recording days. Each dot indicates one recording session. Random, n = 12 days from 7 mice; 40 Hz, n = 12 days from 7 mice; P = 0.12, n.s., LME.

**D.** Sharp-wave ripple duration for SWRs occurring inside the reward zone did not differ significantly following 8 days of 40 Hz (blue) or Random (green) flicker. Random, n = 572 SWRs from 8 mice; 40 Hz, n = 475 SWRs from 8 mice; P = 0.75, n.s., LME.

**E.** Sharp-wave ripple power for SWRs occurring inside the reward zone. Random, n = 572 SWRs from 8 mice; 40 Hz, n = 475 SWRs from 8 mice; P = 0.71, n.s., LME.

**F.** Average SWR abundance in the reward zone did not differ significantly following 8 days of 40 Hz (blue) or Random (green) flicker. SWR abundance was calculated during times in the reward zone where speed was slower than 1 deg/s. Only days with greater than 5 SWRs that met the speed criteria were included. Each dot indicates one recording session. Mean ± SEM across recording days. Random, n = 13 days from 8 mice; 40 Hz, n = 9 days from 6 mice; P = 0.45, n.s., LME.

**G.** Activation probability of place cells in CA1 and CA3 during SWRs occurring inside the reward zone did not differ significantly following 8 days of 40 Hz (blue) or Random (green) flicker. Place cells from sessions with greater than 10 SWRs in the reward zone were included. Random, n = 341 place cells from 7 mice; 40 Hz, n = 426 place cells from 7 mice; P = 0.78, n.s., LME.

**H.** Coactivation probability of all place cell pairs during SWRs occurring inside the reward zone did not differ significantly following 8 days of 40 Hz (blue) or Random (green) flicker. Place cell pairs from sessions with greater than 10 SWRs in the reward zone were included. Random, n = 6519 place cell pairs from 7 mice; 40 Hz, n = 9641 place cell pairs from 7 mice; P = 0.89, n.s., LME.

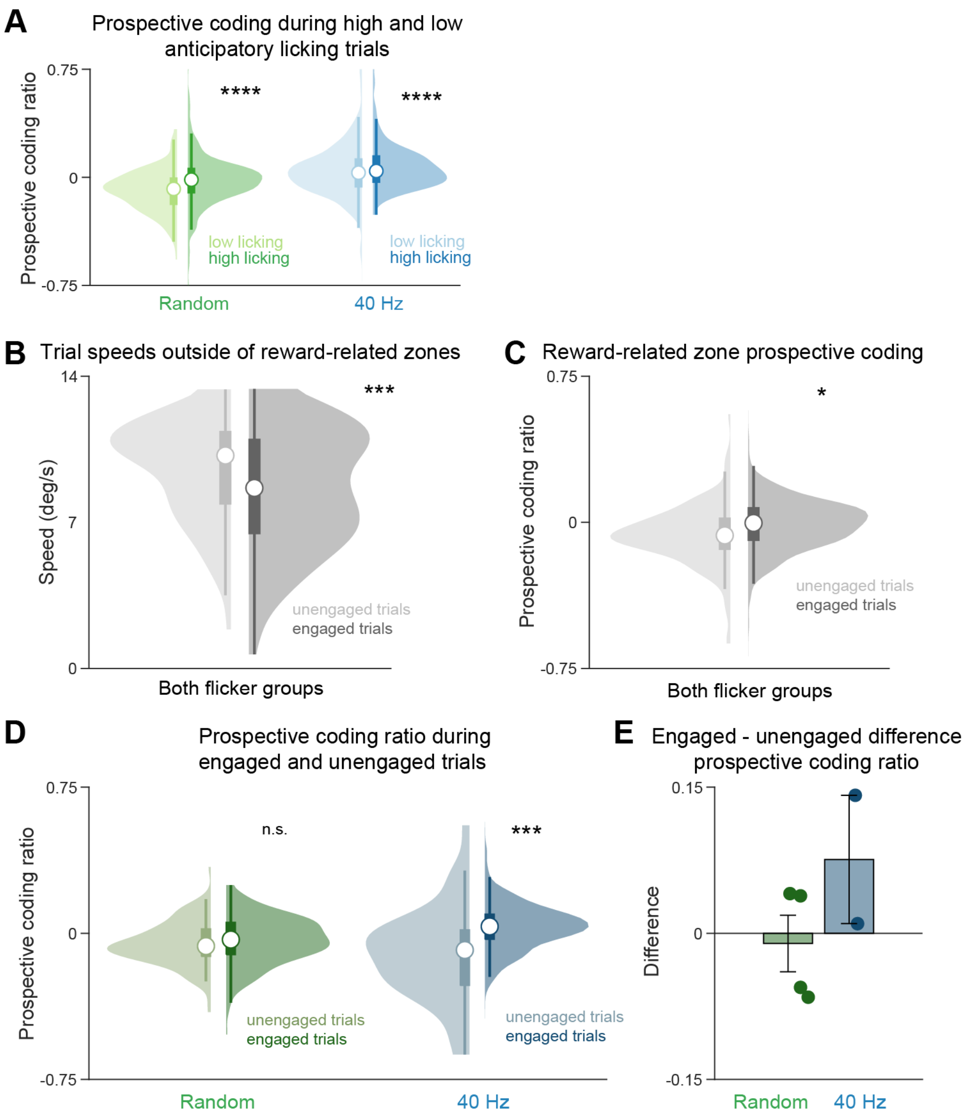


**Supplementary Figure 8. Prospective coding is higher on engaged trials after 40 Hz flicker.**

**A.** After 40 Hz and Random flicker, prospective coding was higher on trials with high anticipatory licking than low anticipatory licking. Prospective coding ratio for trials in the top half (high licks, dark colors) and bottom half (low licks, light colors) of anticipatory lick count. Random, low licks n = 354 trials, high licks = 583 trials, from 8 mice; P = 1.24e-06****, LME. 40 Hz, low licks n = 517 trials, high licks n = 290 trials, from 6 mice; P=2.48e-07****, LME.

**B.** Distributions of trial speeds during unengaged (light grey) and engaged correct (dark grey) trials from Random and 40 Hz flicker groups combined. During unengaged trials, where the animal runs through the environment but does not pause to lick, trial speeds were significantly faster than trials where the animal was engaged in the task and correctly licked in the reward zone. Only data from recording days with both unengaged and engaged trials were included. Unengaged n = 165 trials from 5 mice, engaged n = 455 trials from 5 mice, P = 0.0007***, LME. See Supplementary Table 3 for statistical details.

**C.** Propsective coding ratios did not differ significantly between unengaged (light grey) and engaged correct (dark grey) trials when animals from both flicker groups were considered together. Ratios were calculated in the reward-related areas of the track, as in Figure 3. Unengaged n = 165 trials from 5 mice, engaged n = 455 trials from 5 mice, P = 0.023*, LME.

**D.** Prospective coding ratios were higher after 40 Hz flicker (blue, right) during correct trials when the animal was engaged in the virtual reality task than unengaged trials. After random flicker, prospective coding ratios were not signficaintly different on unengaged and engaged correct trials. Random, unengaged n = 128 trials, engaged 292 trials, from 3 mice, P = 0.49, n.s., LME; 40 Hz, unengaged n = 37 trials, engaged n = 163 trials, from 2 mice, P = 0.0001***, LME.

**E.** Comparing prospective coding on engaged and unengaged trials within each recording day, prospective coding ratios were higher during engaged, correct trials (difference values greater than 0) after 40 Hz flicker. After Random flicker, prospective coding ratio were higher during engaged correct trials on half the days and lower on half the days. Random, n = 4 days from 3 mice, 40 Hz, n = 2 days from 2 mice. Mean +/- SEM. Each dot is a recording day.

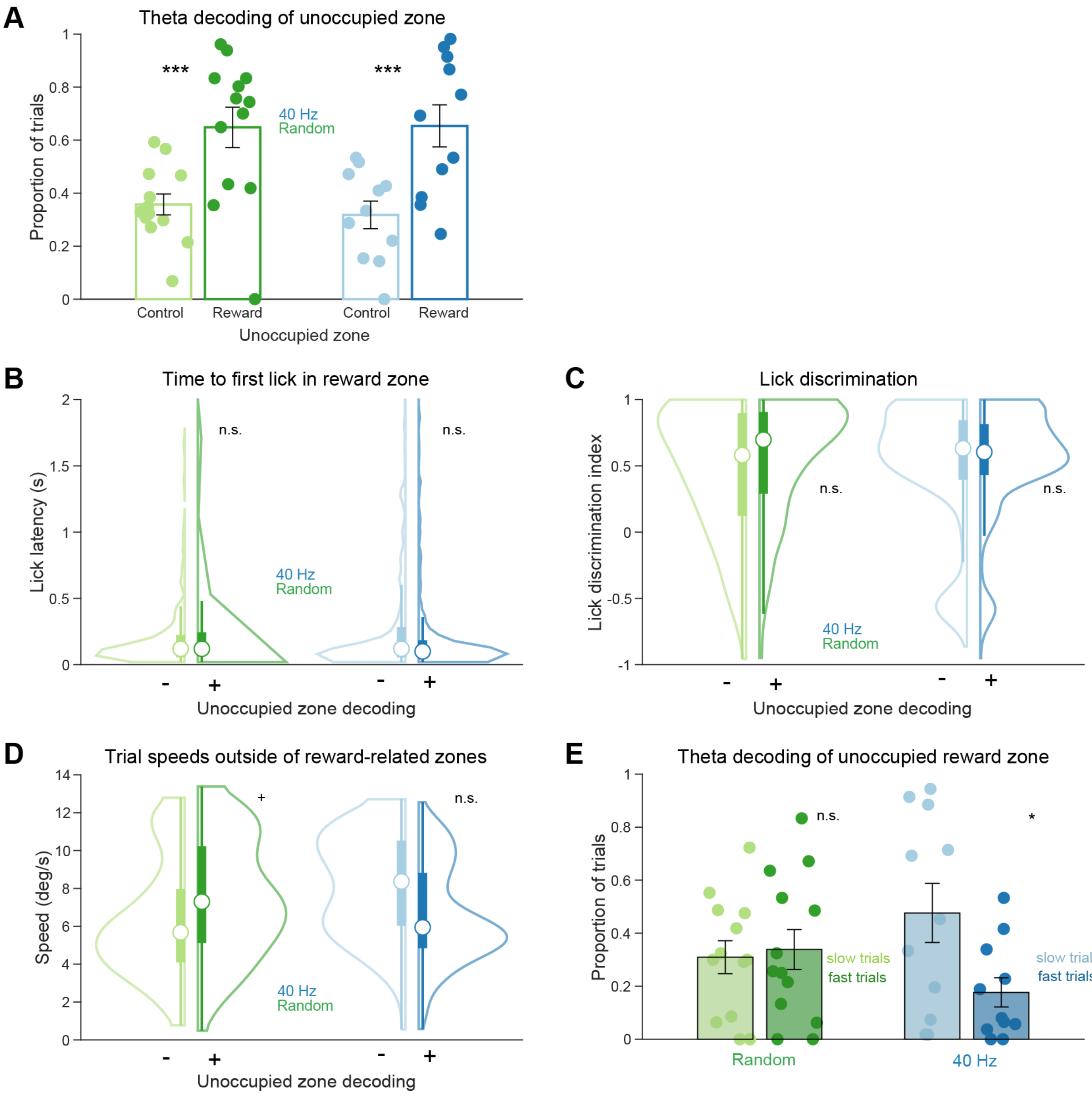


**Supplementary Figure 9. Representation of the unoccupied reward zone during theta.**

- **A.** After both 40 Hz and Random flicker, most trials showed significant decoding of the unoccupied reward zone during theta (left). A larger proportion of trials showed decoding of the unoccupied reward zone, as compared to the unoccupied control zone (right). Random, unoccupied control zone, n = 13 days, unoccupied reward zone, n = 13 days; from 8 mice; P = 0.0014***, LME. 40 Hz, unoccupied control zone, n = 11 days, unoccupied reward zone, n = 11 days; from 6 mice; P = 0.0019***, LME. Mean ± SEM across recording days. Each dot is a recording day. See Supplementary Table 3 for statistical details.
- **B.** Lick latency, the time to the animal's first lick in the reward zones of the track, was not significantly different between trials with or without theta decoding of the unoccupied reward zone. Random, no unoccupied zone decoding, n = 280 trials, unoccupied zone decoding, n = 606 trials; from 8 mice; P = 0.69, n.s., LME. 40 Hz, no unoccupied zone decoding, n = 346 trials, unoccupied zone decoding, n = 474 trials; from 6 mice; P = 0.21, LME.

**C.** Lick discrimination, the proportion of licking occuring in the rewarded areas of the track versus other areas, was not significantly different between trials with or without theta decoding unoccupied reward zone. Random, no unoccupied zone decoding, n = 280 trials, unoccupied zone decoding, n = 606 trials; from 8 mice; P = 0.15, n.s., LME. 40 Hz, no unoccupied zone decoding, n = 346 trials, unoccupied zone decoding, n = 474 trials; from 6 mice; P = 0.21, LME.

**D.** Trial speed, calculated outside of the reward-related areas of the track, was not significantly different between trials with or without theta decoding of the unoccupied reward zone. Random, no unoccupied zone decoding, n = 280 trials, unoccupied zone decoding, n = 606 trials; from 8 mice; P = 0.06+, LME. 40 Hz, no unoccupied zone decoding, n = 346 trials, unoccupied zone decoding, n = 474 trials; from 6 mice; P = 0.87, LME.

**E.** Proportion of slow or fast trials that contain theta decoding of the unoccupied reward zone. After 40 Hz flicker, slow trials contain a significantly larger proportion of trials that have significant decoding of the other reward zone. Random, proportion of slow trials, n = 13 days, proportion of fast trials, n = 13 days; from 8 mice; P = 0.98, n.s., LME. 40 Hz, proportion of slow trials, n = 11 days, proportion of fast trials n = 11 days; from 6 mice; P = 0.028*, LME. Mean $\pm$ SEM across recording days. Each dot is a recording day.

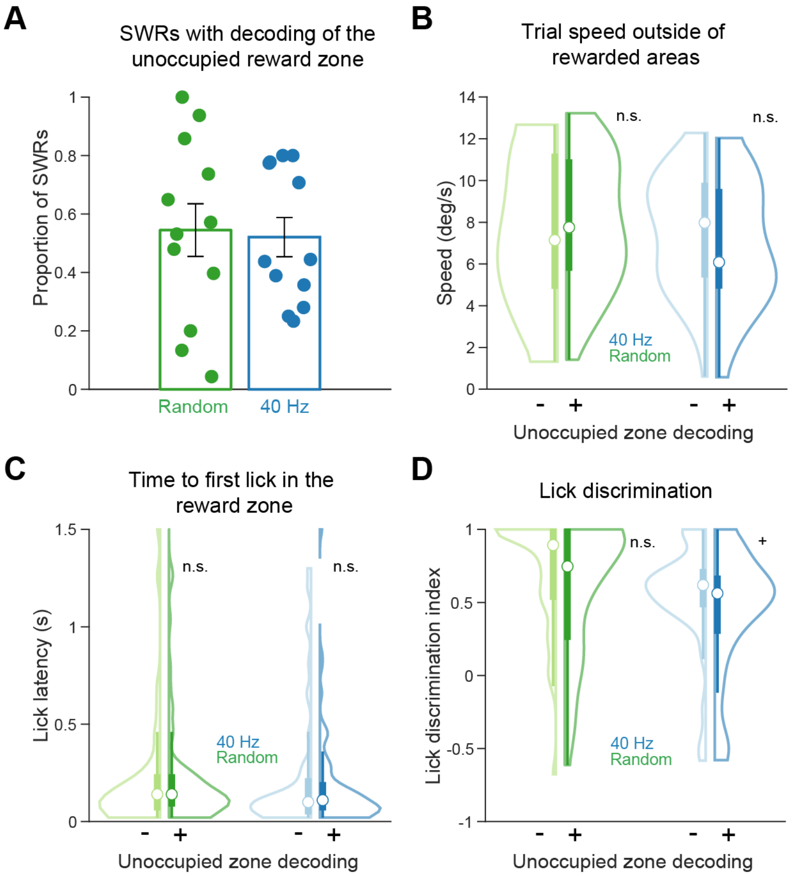


**Supplementary Figure 10. Representation of the unoccupied reward zone during SWRs.**

- **A.** Proportion of SWRs that contain significant position information representing the unoccupied reward zone. Only SWRs occurring in the reward zones were included for analysis. Random, n = 12 days from 7 mice; 40 Hz, n = 12 days from 7 mice. Mean ± SEM across recording days. Each dot is a recording day. See Supplementary Table 3 for statistical details.
- **B.** Trial speed, calculated outside of the reward-related areas of the track, was not significantly different between trials with (+) or without (-) decoding of the unoccupied reward zone during SWRs. Random, no unoccupied zone decoding (-), n = 42 trials, unoccupied zone decoding (+), n = 94 trials; P = 0.40, n.s., LME. 40 Hz, no unoccupied zone decoding (-), n = 84 trials, unoccupied zone decoding (+), n = 76 trials; P = 0.12, n.s., LME.
- **C.** Lick latency, the time to the animal's first lick in the reward zones of the track, was not significantly different between trials with or without decoding of the unoccupied reward zone during SWRs. Random trials, no unoccupied zone decoding, 1 data point above 1.5, and Random trials with unoccupied zone decoding, 2 data points above 1.5, are excluded for visualization purposes. Random, no unoccupied zone decoding (-), n = 42 trials, unoccupied zone decoding (+), n = 94 trials; P = 0.68, n.s., LME. 40 Hz, no unoccupied zone decoding (-), n = 84 trials, unoccupied zone decoding (+), n = 76 trials; P = 0.25, n.s., LME.
- **D.** Lick discrimination, the proportion of licking occuring in the rewarded areas of the track versus other areas, was not significantly different between trials with or without decoding of unoccupied reward zone during SWRs. Random, no unoccupied zone decoding (-), n = 42 trials, unoccupied zone decoding (+), n = 94 trials; P = 0.41, n.s., LME. 40 Hz, no unoccupied zone decoding (-), n = 84 trials, unoccupied zone decoding (+), n = 76 trials; P = 0.07+, LME.

| AnimalID | Flicker Group | Recorded Cells | CA1 cells | CA3 cells | Putative Pyramidal Cells | Putative Interneurons | Place cells | # Trials | # SWRs |
|---|---|---|---|---|---|---|---|---|---|
| 28 | 40 Hz | 171 | 103 | 68 | 128 | 9 | 102 | 145 | 41 |
| 29 | Random | 182 | 91 | 91 | 147 | 12 | 107 | 154 | 133 |
| 30 | 40 Hz | 140 | 79 | 61 | 105 | 12 | 81 | 48 | 1224 |
| 31 | Random | 92 | 22 | 70 | 49 | 10 | 21 | 107 | 725 |
| 32 | 40 Hz | 199 | 106 | 93 | 146 | 17 | 98 | 80 | 294 |
| 35 | 40 Hz | 93 | 38 | 55 | 75 | 4 | 49 | 20 | 131 |
| 36 | Random | 164 | 80 | 84 | 128 | 9 | 69 | 142 | 130 |
| 37 | Random | 188 | 86 | 102 | 117 | 20 | 66 | 115 | 318 |
| 39 | 40 Hz | 117 | 27 | 90 | 78 | 7 | 42 | 175 | 52 |
| 40 | Random | 139 | 88 | 51 | 109 | 7 | 46 | 140 | 76 |
| 41 | 40 Hz | 218 | 78 | 140 | 170 | 5 | 79 | 25 | 156 |
| 42 | Random | 180 | 95 | 85 | 127 | 9 | 52 | 111 | 110 |
| 45 | 40 Hz | 196 | 118 | 78 | 150 | 11 | 74 | 115 | 12 |
| 46 | Random | 136 | 76 | 60 | 84 | 10 | 46 | 133 | 322 |
| 47 | 40 Hz | 109 | 63 | 46 | 82 | 7 | 58 | 292 | 239 |
| 48 | Random | 141 | 76 | 65 | 104 | 5 | 64 | 95 | 239 |

**Supplementary Table 1.** Recording information.

| Figure | Panel | Analysis | Linear Mixed-Effects | | | Group | N | Unit | # Mice | Mean | SEM | Median | 1st Quartile | 3rd Quartile |
|---|---|---|---|---|---|---|---|---|---|---|---|---|---|---|
| | | | P-value | Notation | F-statistic | | | | | | | | | |
| 2 | C | PYR slow gamma PPC flicker periods | 0.433 | ns | $F_{1,13.89} = 0.65$ | random | 320 | cells | 8 | 5.48E-05 | 3.06E-06 | 2.96E-05 | 1.65E-06 | 7.61E-05 |
| | | | | | | 40 Hz | 387 | cells | 8 | 8.91E-05 | 4.53E-06 | 6.17E-05 | 1.63E-05 | 1.18E-04 |
| | C | PYR slow gamma PPC VR task | 0.024 | * | $F_{1,13.90} = 6.42$ | random | 347 | cells | 8 | 3.99E-05 | 2.14E-06 | 2.26E-05 | 3.83E-06 | 5.30E-05 |
| | | | | | | 40 Hz | 335 | cells | 8 | 1.13E-04 | 6.19E-06 | 6.92E-05 | 1.74E-05 | 1.44E-04 |
| | D | IN slow gamma PPC flicker periods | 0.097 | + | $F_{1,11.76} = 3.26$ | random | 46 | cells | 8 | 8.46E-05 | 1.25E-05 | 5.37E-05 | 1.22E-05 | 1.23E-04 |
| | | | | | | 40 Hz | 50 | cells | 8 | 1.79E-04 | 2.53E-05 | 1.08E-04 | 4.21E-05 | 1.96E-04 |
| | D | IN slow gamma PPC VR task | 0.021 | * | $F_{1,8.88} = 7.81$ | random | 49 | cells | 8 | 8.83E-05 | 1.26E-05 | 5.74E-05 | 2.43E-05 | 1.25E-04 |
| | | | | | | 40 Hz | 45 | cells | 8 | 2.03E-04 | 3.03E-05 | 1.14E-04 | 6.10E-05 | 2.57E-04 |
| 3 | E | theta prospective coding ratio | 0.003 | ** | $F_{1,10.63} = 15.15$ | random | 13 | days | 8 | -0.033 | 0.017 | -0.019 | -0.062 | -0.004 |
| | | | | | | 40 Hz | 10 | days | 6 | 0.052 | 0.016 | 0.054 | 0.025 | 0.090 |
| | F | prospective coding slow trials | 0.005 | ** | $F_{1,11.08} = 12.57$ | random | 480 | trials | 8 | -0.039 | 0.010 | -0.030 | -0.122 | 0.048 |
| | | | | | | 40 Hz | 389 | trials | 6 | 0.050 | 0.009 | 0.035 | -0.059 | 0.127 |
| | F | prospective coding fast trials | 0.0007 | *** | $F_{1,7.99} = 28.56$ | random | 454 | trials | 8 | -0.061 | 0.008 | -0.062 | -0.146 | 0.032 |
| | | | | | | 40 Hz | 418 | trials | 6 | 0.064 | 0.009 | 0.043 | -0.042 | 0.147 |
| | H | reward-related place cells | 0.783 | ns | $F_{1,11.80} = 0.079$ | random | 13 | days | 8 | 0.169 | 0.027 | 0.214 | 0.062 | 0.236 |
| | | | | | | 40 Hz | 10 | days | 6 | 0.166 | 0.020 | 0.160 | 0.107 | 0.219 |
| 4 | C | SWR prospective coding ratio | 0.047 | * | $F_{1,11.30} = 4.95$ | random | 507 | SWRs | 7 | 0.047 | 0.012 | 0.041 | -0.109 | 0.208 |
| | | | | | | 40 Hz | 378 | SWRs | 7 | 0.200 | 0.016 | 0.212 | -0.001 | 0.433 |
| | E | reward zone prospective SWRs | 0.033 | * | $F_{1,11.22} = 5.88$ | random | 12 | days | 7 | 0.387 | 0.061 | 0.405 | 0.203 | 0.568 |
| | | | | | | 40 Hz | 12 | days | 7 | 0.606 | 0.056 | 0.633 | 0.5 | 0.75 |
| 5 | C | trial speed slow trials | 0.214 | ns | $F_{1,11.77} = 1.72$ | random | 483 | trials | 8 | 4.495 | 0.064 | 4.692 | 3.538 | 5.684 |
| | | | | | | 40 Hz | 389 | trials | 6 | 5.028 | 0.055 | 5.243 | 4.449 | 5.804 |
| | C | trial speed fast trials | 0.993 | ns | $F_{1,12.12} = 9e\text{-}05$ | random | 454 | trials | 8 | 9.600 | 0.093 | 9.516 | 7.693 | 11.428 |
| | | | | | | 40 Hz | 418 | trials | 6 | 9.489 | 0.080 | 9.627 | 8.227 | 10.828 |
| | E | prospective coding random | 0.009 | ** | $F_{1,929.14} = 6.93$ | slow trials | 483 | trials | 8 | 0.047 | 0.008 | 0.013 | -0.043 | 0.113 |
| | | | | | | fast trials | 454 | trials | 8 | 0.051 | 0.007 | 0.041 | -0.037 | 0.132 |
| | E | prospective coding 40 Hz | 1.10E-09 | **** | $F_{1,490.72} = 38.58$ | slow trials | 389 | trials | 6 | -0.007 | 0.007 | -0.012 | -0.083 | 0.066 |
| | | | | | | fast trials | 418 | trials | 6 | 0.109 | 0.008 | 0.092 | 0.003 | 0.209 |
| | F | time to next reward zone random | <2e-16 | **** | $F_{1,728.93} = 185.27$ | slow trials | 386 | trials | 8 | 35.16 | 0.696 | 31.94 | 28.06 | 37.56 |
| | | | | | | fast trials | 417 | trials | 8 | 18.55 | 0.205 | 17.46 | 15.5 | 20.92 |
| | F | time to next reward zone 40 Hz | <2e-16 | **** | $F_{1,415.98} = 173.74$ | slow trials | 475 | trials | 6 | 43.93 | 1.29 | 37.3 | 29.92 | 47.8 |
| | | | | | | fast trials | 454 | trials | 6 | 19.73 | 0.26 | 18.11 | 15.46 | 22.8 |

**Supplementary Table 2.** Statistical information for main figures. Pyramidal cells (PYR); Interneurons (IN).

| Figure | Panel | Analysis | P-value | Notation | F-statistic | Group | N | Unit | # Mice | Mean | SEM | Median | 1st Quartile | 3rd Quartile |
|---|---|---|---|---|---|---|---|---|---|---|---|---|---|---|
| S1 | G | speed | 0.51 | ns | $F_{1,13.96} = 0.46$ | random | 15 | days | 8 | 3.784 | 0.347 | 3.764 | 2.818 | 4.818 |
| | | | | | | 40 Hz | 15 | days | 8 | 3.455 | 0.372 | 2.746 | 2.638 | 4.975 |
| | I | lick latency | 0.74 | ns | $F_{1,13.86} = 0.12$ | random | 15 | days | 8 | 0.464 | 0.181 | 0.251 | 0.149 | 0.361 |
| | | | | | | 40 Hz | 15 | days | 8 | 0.578 | 0.207 | 0.209 | 0.158 | 0.555 |
| S4 | C | PYR theta PPC flicker periods | 0.19 | ns | $F_{1,13.19} = 1.91$ | random | 320 | cells | 8 | 2.07E-03 | 1.16E-04 | 2.00E-03 | 1.42E-03 | 2.58E-03 |
| | | | | | | 40 Hz | 387 | cells | 8 | 1.62E-03 | 8.21E-05 | 1.61E-03 | 8.18E-04 | 2.29E-03 |
| | C | PYR theta PPC VR task | 0.29 | ns | $F_{1,14.00} = 1.21$ | random | 347 | cells | 8 | 1.00E-03 | 5.39E-05 | 7.41E-04 | 3.50E-04 | 1.36E-03 |
| | | | | | | 40 Hz | 335 | cells | 8 | 1.66E-03 | 9.08E-05 | 1.13E-03 | 5.86E-04 | 2.00E-03 |
| | D | IN theta PPC flicker periods | 0.36 | ns | $F_{1,8.93} = 0.93$ | random | 46 | cells | 8 | 2.52E-03 | 3.72E-04 | 2.47E-03 | 1.24E-03 | 3.61E-03 |
| | | | | | | 40 Hz | 50 | cells | 8 | 2.86E-03 | 4.04E-04 | 2.63E-03 | 1.87E-03 | 3.54E-03 |
| | D | IN theta PPC VR task | 0.12 | ns | $F_{1,13.23} = 2.73$ | random | 49 | cells | 8 | 1.73E-03 | 2.47E-04 | 0.001384 | 7.06E-04 | 2.02E-03 |
| | | | | | | 40 Hz | 50 | cells | 8 | 2.68E-03 | 3.79E-04 | 2.10E-03 | 1.30E-03 | 3.18E-03 |
| | E | PYR medium 40 Hz PPC flicker periods | 0.90 | ns | $F_{1,13.29} = 0.015$ | random | 320 | cells | 8 | 1.40E-05 | 7.85E-07 | 4.89E-06 | -5.39E-06 | 2.43E-05 |
| | | | | | | 40 Hz | 387 | cells | 8 | 1.64E-05 | 8.35E-07 | 7.74E-06 | -3.00E-07 | 2.64E-05 |
| | E | PYR medium 40 Hz PPC VR task | 0.32 | ns | $F_{1,12.99} = 1.09$ | random | 347 | cells | 8 | 8.57E-06 | 4.60E-07 | 1.61E-07 | -5.35E-06 | 1.79E-05 |
| | | | | | | 40 Hz | 335 | cells | 8 | 1.53E-05 | 8.37E-07 | 4.25E-06 | -4.60E-06 | 2.27E-05 |
| | F | IN medium 40 Hz PPC flicker periods | 0.21 | ns | $F_{1,11.52} = 1.75$ | random | 46 | cells | 8 | 2.67E-05 | 3.93E-06 | 1.39E-05 | 5.62E-06 | 3.70E-05 |
| | | | | | | 40 Hz | 50 | cells | 8 | 4.52E-05 | 6.39E-06 | 2.31E-05 | 8.28E-06 | 5.20E-05 |
| | F | IN medium 40 Hz PPC VR task | 0.11 | ns | $F_{1,8.88} = 3.10$ | random | 49 | cells | 8 | 2.83E-05 | 4.05E-06 | 1.87E-05 | 6.87E-06 | 4.17E-05 |
| | | | | | | 40 Hz | 45 | cells | 8 | 4.68E-05 | 6.98E-06 | 2.28E-05 | 8.68E-06 | 6.96E-05 |
| S6 | B | spatial information | 0.93 | ns | $F_{1,13.98} = 0.01$ | random | 471 | place cells | 8 | 0.253 | 0.010 | 0.197 | 0.107 | 0.318 |
| | | | | | | 40 Hz | 583 | place cells | 8 | 0.244 | 0.010 | 0.177 | 0.081 | 0.318 |
| | C | sparsity | 0.69 | ns | $F_{1,13.98} = 0.17$ | random | 471 | place cells | 8 | 0.183 | 0.008 | 0.122 | 0.048 | 0.266 |
| | | | | | | 40 Hz | 583 | place cells | 8 | 0.190 | 0.008 | 0.126 | 0.048 | 0.282 |
| | D | peak position | 0.68 | ns | $F_{1,13.58} = 0.18$ | random | 471 | place cells | 8 | 9.013 | 2.369 | 6.000 | -30.000 | 54.000 |
| | | | | | | 40 Hz | 583 | place cells | 8 | 11.027 | 2.115 | 12.000 | -27.000 | 51.000 |
| | E | peak firing rate | 0.86 | ns | $F_{1,13.58} = 0.03$ | random | 471 | place cells | 8 | 3.731 | 0.134 | 2.772 | 1.523 | 4.950 |
| | | | | | | 40 Hz | 583 | place cells | 8 | 3.646 | 0.115 | 2.748 | 1.533 | 5.165 |
| | F | mean firing rate | 0.85 | ns | $F_{1,13.68} = 0.04$ | random | 471 | place cells | 8 | 2.127 | 0.095 | 1.367 | 0.652 | 2.854 |
| | | | | | | 40 Hz | 583 | place cells | 8 | 2.103 | 0.085 | 1.293 | 0.640 | 2.871 |
| | G | theta sequence quadrant ratio | 0.3702 | ns | $F_{1,10.63} = 0.88$ | random | 13 | days | 8 | 0.017 | 0.003 | 0.013 | 0.009 | 0.026 |
| | | | | | | 40 Hz | 10 | days | 6 | 0.023 | 0.007 | 0.020 | 0.009 | 0.025 |
| | I | prospective coding ratio | 0.0007 | *** | $F_{1,11.92} = 20.82$ | random | 934 | trials | 8 | -0.049 | 0.006 | -0.048 | -0.134 | 0.045 |
| | | | | | | 40 Hz | 807 | trials | 6 | 0.057 | 0.006 | 0.037 | -0.049 | 0.137 |
| | J | RRZ vs. control prospective coding ratio | 0.0089 | ** | $F_{1,1856.02} = 6.85$ | random; reward zone | 934 | trials | 8 | -0.049 | 0.006 | -0.048 | -0.134 | 0.045 |
| | | | | | | random; control zone | 931 | trials | 8 | -0.029 | 0.005 | -0.027 | -0.123 | 0.062 |
| | J | RRZ vs. control prospective coding ratio | 1.4E-13 | **** | $F_{1,1599.14} = 55.70$ | 40 Hz; reward zone | 807 | trials | 6 | 0.057 | 0.006 | 0.037 | -0.049 | 0.137 |
| | | | | | | 40 Hz; control zone | 799 | trials | 6 | -0.028 | 0.009 | -0.029 | -0.149 | 0.101 |
| S7 | A | all SWR count | 0.46 | ns | $F_{1,11.76} = 0.59$ | random | 12 | days | 7 | 95.42 | 19.43 | 95.50 | 31.50 | 126.50 |
| | | | | | | 40 Hz | 12 | days | 7 | 165.42 | 69.82 | 45.50 | 20.50 | 229.00 |
| | B | SWR prospective coding ratio | 0.21 | ns | $F_{1,11.87} = 1.77$ | random | 1145 | SWRs | 7 | 0.082 | 0.008 | 0.073 | -0.095 | 0.261 |
| | | | | | | 40 Hz | 1989 | SWRs | 7 | 0.176 | 0.006 | 0.188 | 0.013 | 0.355 |
| | C | prospective SWRs | 0.12 | ns | $F_{1,11.27} = 2.88$ | random | 12 | days | 7 | 0.417 | 0.066 | 0.408 | 0.230 | 0.589 |
| | | | | | | 40 Hz | 12 | days | 7 | 0.587 | 0.060 | 0.629 | 0.461 | 0.724 |
| | D | SWR duration | 0.75 | ns | $F_{1,23.55} = 0.10$ | random | 572 | SWRs | 8 | 0.101 | 0.002 | 0.093 | 0.072 | 0.122 |
| | | | | | | 40 Hz | 475 | SWRs | 8 | 0.101 | 0.002 | 0.091 | 0.071 | 0.123 |
| | E | SWR power | 0.41 | ns | $F_{1,24.17} = 0.71$ | random | 572 | SWRs | 8 | 5.101 | 0.092 | 4.307 | 3.527 | 5.843 |
| | | | | | | 40 Hz | 475 | SWRs | 8 | 5.114 | 0.128 | 4.208 | 3.391 | 5.684 |
| | F | SWR abundance | 0.45 | ns | $F_{1,10.72} = 0.62$ | random | 13 | days | 8 | 0.027 | 0.004 | 0.023 | 0.019 | 0.041 |
| | | | | | | 40 Hz | 9 | days | 6 | 0.034 | 0.008 | 0.027 | 0.020 | 0.048 |
| | G | activation probability | 0.78 | ns | $F_{1,11.78} = 0.084$ | random | 341 | place cells | 7 | 0.388 | 0.014 | 0.333 | 0.192 | 0.547 |
| | | | | | | 40 Hz | 426 | place cells | 7 | 0.370 | 0.012 | 0.333 | 0.146 | 0.559 |
| | H | coactivation probability | 0.89 | ns | $F_{1,12.00} = 0.018$ | random | 6519 | place cell pairs | 7 | 0.182 | 0.002 | 0.123 | 0.053 | 0.250 |
| | | | | | | 40 Hz | 9641 | place cell pairs | 7 | 0.180 | 0.002 | 0.111 | 0.049 | 0.250 |
| S8 | A | propspective coding ratio by anticipatory licking | 1.2E-06 | **** | $F_{1,920.43} = 23.83$ | random; low licking | 346 | trials | 8 | -0.091 | 0.008 | -0.080 | -0.187 | -0.003 |
| | | | | | | random; high licking | 588 | trials | 8 | -0.025 | 0.009 | -0.016 | -0.108 | 0.063 |
| | A | propspective coding ratio by anticipatory licking | 2.5E-07 | **** | $F_{1,729.03} = 27.13$ | 40 Hz; low licking | 517 | trials | 6 | 0.048 | 0.009 | 0.033 | -0.066 | 0.129 |
| | | | | | | 40 Hz; high licking | 290 | trials | 6 | 0.074 | 0.010 | 0.044 | -0.034 | 0.150 |
| | B | engaged vs. unengaged trial speeds | 0.0007 | *** | $F_{1,615.97} = 11.67$ | unengaged trials | 165 | trials | 5 | 9.492 | 0.187 | 10.201 | 7.879 | 11.350 |
| | | | | | | engaged trials | 455 | trials | 5 | 8.517 | 0.135 | 8.644 | 6.457 | 10.975 |
| | C | engaged vs. unengaged prospective coding ratio | 0.02 | * | $F_{1,615.04} = 5.15$ | unengaged trials | 165 | trials | 5 | -0.066 | 0.012 | -0.067 | -0.139 | 0.022 |
| | | | | | | engaged trials | 455 | trials | 5 | -0.012 | 0.007 | -0.004 | -0.093 | 0.075 |
| | D | engaged vs. unengaged prospective coding ratio | 0.49 | ns | $F_{1,416.84} = 0.48$ | random; unengaged trials | 128 | trials | 3 | -0.055 | 0.010 | -0.065 | -0.118 | 0.023 |
| | | | | | | random; engaged trials | 292 | trials | 3 | -0.036 | 0.007 | -0.032 | -0.109 | 0.057 |
| | D | engaged vs. unengaged prospective coding ratio | 0.0001 | *** | $F_{1,197.95} = 15.68$ | 40 Hz; unengaged trials | 37 | trials | 2 | -0.105 | 0.040 | -0.086 | -0.267 | 0.018 |
| | | | | | | 40 Hz; engaged trials | 163 | trials | 2 | 0.032 | 0.012 | 0.036 | -0.031 | 0.099 |
| S9 | A | unoccupied zone decoding proportion of trials | 0.0001 | *** | $F_{1,17.04} = 23.97$ | random; reward zone | 13 | days | 8 | 0.357 | 0.040 | 0.329 | 0.291 | 0.468 |
| | | | | | | random; control zone | 13 | days | 8 | 0.648 | 0.076 | 0.744 | 0.430 | 0.833 |
| | A | unoccupied zone decoding proportion of trials | 0.0002 | *** | $F_{1,15.02} = 24.03$ | 40 Hz; reward zone | 11 | days | 6 | 0.318 | 0.052 | 0.333 | 0.170 | 0.460 |
| | | | | | | 40 Hz; control zone | 11 | days | 6 | 0.653 | 0.079 | 0.692 | 0.411 | 0.902 |
| | B | unoccupied zone decoding lick latency | 0.69 | ns | $F_{1,505.48} = 0.16$ | random; no unoccupied zone decoding | 280 | trials | 8 | 0.219 | 0.024 | 0.120 | 0.060 | 0.220 |
| | | | | | | random; unoccupied zone decoding | 606 | trials | 8 | 0.362 | 0.059 | 0.120 | 0.080 | 0.240 |
| | B | unoccupied zone decoding lick latency | 0.21 | ns | $F_{1,817.53} = 1.55$ | 40 Hz; no unoccupied zone decoding | 346 | trials | 6 | 0.298 | 0.026 | 0.120 | 0.060 | 0.280 |
| | | | | | | 40 Hz; unoccupied zone decoding | 474 | trials | 6 | 0.211 | 0.017 | 0.100 | 0.060 | 0.180 |
| | C | unoccupied zone decoding lick discrimination | 0.15 | ns | $F_{1,880.41} = 2.06$ | random; no unoccupied zone decoding | 280 | trials | 8 | 0.468 | 0.028 | 0.581 | 0.127 | 0.894 |
| | | | | | | random; unoccupied zone decoding | 606 | trials | 8 | 0.553 | 0.018 | 0.698 | 0.295 | 0.903 |
| | C | unoccupied zone decoding lick discrimination | 0.21 | ns | $F_{1,817.49} = 1.58$ | 40 Hz; no unoccupied zone decoding | 346 | trials | 6 | 0.504 | 0.026 | 0.632 | 0.399 | 0.840 |
| | | | | | | 40 Hz; unoccupied zone decoding | 474 | trials | 6 | 0.547 | 0.019 | 0.604 | 0.434 | 0.810 |
| | D | unoccupied zone decoding trial speeds | 0.07 | + | $F_{1,880.96} = 3.39$ | random; no unoccupied zone decoding | 280 | trials | 8 | 6.250 | 0.179 | 5.681 | 4.125 | 7.949 |
| | | | | | | random; unoccupied zone decoding | 606 | trials | 8 | 7.488 | 0.125 | 7.301 | 5.155 | 10.194 |
| | D | unoccupied zone decoding trial speeds | 0.87 | ns | $F_{1,814.94} = 0.026$ | 40 Hz; no unoccupied zone decoding | 346 | trials | 6 | 8.139 | 0.142 | 8.368 | 6.060 | 10.502 |
| | | | | | | 40 Hz; unoccupied zone decoding | 474 | trials | 6 | 6.585 | 0.119 | 5.945 | 4.871 | 8.793 |
| | E | unoccupied zone decoding by median trial speed | 0.77 | ns | $F_{1,17.49} = 0.09$ | Random; slow trials | 13 | days | 8 | 0.309 | 0.062 | 0.300 | 0.080 | 0.479 |
| | | | | | | Random; fast trials | 13 | days | 8 | 0.339 | 0.075 | 0.256 | 0.116 | 0.559 |
| | E | unoccupied zone decoding by median trial speed | 0.03 | * | $F_{1,15.19} = 5.82$ | 40 Hz; slow trials | 11 | days | 6 | 0.476 | 0.111 | 0.453 | 0.104 | 0.843 |
| | | | | | | 40 Hz; fast trials | 11 | days | 6 | 0.177 | 0.055 | 0.080 | 0.042 | 0.311 |
| S10 | B | SWR unoccupied zone decoding trial speed | 0.40 | ns | $F_{1,128.91} = 0.72$ | random; no unoccupied zone decoding | 42 | trials | 7 | 7.572 | 0.543 | 7.143 | 4.833 | 11.256 |
| | | | | | | random; unoccupied zone decoding | 94 | trials | 7 | 7.954 | 0.314 | 7.756 | 5.702 | 10.987 |
| | B | SWR unoccupied zone decoding trial speed | 0.12 | ns | $F_{1,154.59} = 2.51$ | 40 Hz; no unoccupied zone decoding | 84 | trials | 7 | 7.594 | 0.292 | 7.979 | 5.379 | 9.861 |
| | | | | | | 40 Hz; unoccupied zone decoding | 76 | trials | 7 | 6.828 | 0.338 | 6.084 | 4.844 | 9.567 |
| | C | SWR unoccupied zone decoding lick latency | 0.68 | ns | $F_{1,134.0} = 0.17$ | random; no unoccupied zone decoding | 42 | trials | 7 | 0.285 | 0.079 | 0.140 | 0.060 | 0.240 |
| | | | | | | random; unoccupied zone decoding | 94 | trials | 7 | 0.256 | 0.039 | 0.140 | 0.080 | 0.240 |
| | C | SWR unoccupied zone decoding lick latency | 0.25 | ns | $F_{1,155.79} = 1.33$ | 40 Hz; no unoccupied zone decoding | 84 | trials | 7 | 0.234 | 0.035 | 0.100 | 0.040 | 0.220 |
| | | | | | | 40 Hz; unoccupied zone decoding | 76 | trials | 7 | 0.204 | 0.033 | 0.110 | 0.060 | 0.200 |
| | D | SWR unoccupied zone decoding lick | 0.41 | ns | $F_{1,129.26} = 0.69$ | random; no unoccupied zone decoding | 42 | trials | 7 | 0.714 | 0.060 | 0.893 | 0.522 | 1.000 |
| | | | | | | random; unoccupied zone decoding | 94 | trials | 7 | 0.592 | 0.047 | 0.747 | 0.247 | 1.000 |
| | D | SWR unoccupied zone decoding lick | 0.07 | + | $F_{1,155.86} = 3.34$ | 40 Hz; no unoccupied zone decoding | 84 | trials | 7 | 0.550 | 0.039 | 0.619 | 0.471 | 0.726 |
| | | | | | | 40 Hz; unoccupied zone decoding | 76 | trials | 7 | 0.434 | 0.049 | 0.562 | 0.289 | 0.680 |

**Supplementary Table 3.** Statistical information for supplemental figures.

| Figure | Panel | Analysis | Significant Frequency Points ($p < 0.05$, unpaired t-test without correction) |
|---|---|---|---|
| **S3** | **A** | WPLI VR | 14.1602, 14.6484, 15.1367, 15.625, 16.1133, 16.6016, 17.0898, 17.5781, 18.0664, 18.5547, 19.043, 23.9258, 26.8555, 27.3438, 28.8086, 29.2969, 29.7852, 30.2734, 30.7617, 31.7383, 32.7148, 34.1797, 34.668, 35.1562, 36.1328, 36.6211 |
| **S3** | **A** | WPLI Flicker periods | 10.2539, 10.7422, 11.2305, 11.7188, 12.207, 12.6953, 15.1367, 15.625, 16.1133, 16.6016, 17.0898, 17.5781, 18.0664, 18.5547, 19.043, 19.5312, 20.0195, 20.5078, 20.9961, 21.9727, 22.4609, 24.9023, 35.6445, 39.5508, 40.0391, 119.6289 |
| **S3** | **B** | CA1 PSD VR | 6.8359, 24.9023, 26.8555, 27.3438 |
| **S3** | **B** | CA1 PSD Flicker periods | 18.5547, 21.9727, 22.4609, 24.9023, 25.3906, 26.3672, 26.8555, 39.5508, 40.0391, 79.5898, 80.0781, 119.6289 |
| **S3** | **C** | CA3 PSD VR | 56.1523, 59.5703, 60.0586, 60.5469, 61.5234, 62.0117, 63.4766, 63.9648, 64.4531, 64.9414, 65.4297, 65.918, 66.8945, 67.3828, 67.8711, 68.3594, 68.8477, 69.8242, 70.3125, 70.8008, 71.2891, 71.7773, 72.7539, 74.2188, 74.707, 75.6836, 77.1484, 79.1016 |
| **S3** | **C** | CA3 PSD Flicker periods | 5.8594, 13.6719, 14.1602, 39.5508, 40.0391, 52.7344, 55.1758, 56.1523, 56.6406, 57.1289, 57.6172, 58.1055, 58.5938, 59.082, 59.5703, 60.0586, 60.5469, 61.0352, 61.5234, 62.0117, 62.5, 62.9883, 63.4766, 63.9648, 64.4531, 64.9414, 65.4297, 65.918, 66.4062, 66.8945, 67.3828, 67.8711, 68.3594, 68.8477, 69.3359, 69.8242, 70.3125, 70.8008, 71.2891, 71.7773, 72.2656, 72.7539, 73.2422, 73.7305, 74.2188, 74.707, 75.1953, 75.6836, 76.1719, 76.6602, 77.1484, 77.6367, 78.125, 78.6133, 79.1016, 79.5898, 80.0781, 81.0547, 81.543, 82.0312, 82.5195, 83.0078, 83.4961, 83.9844, 84.4727, 84.9609, 85.4492, 85.9375, 86.4258, 86.9141, 87.4023, 87.8906, 88.3789, 88.8672, 89.3555, 89.8438, 90.332, 90.8203, 91.3086, 91.7969, 92.2852, 92.7734, 93.2617, 94.2383, 94.7266, 95.2148, 95.7031, 96.1914, 96.6797, 97.168, 99.6094, 100.0977, 101.5625, 102.0508, 110.3516 |

**Supplementary Table 4**. Statistical information for Supplementary Figure 3. Significant frequency points (indicated by * in Supplementary Figure 3) are listed here. Weighted phase-lag index (WPLI); power spectral density (PSD).

| Figure | Analysis | Cells per animal (min-max) | |
|---|---|---|---|
| | | **Random** | **40 Hz** |
| 2C, left | CA3 Pyramidal Cell slow gamma PPC during flicker periods | 8 - 67 | 28 - 99 |
| 2C, right | CA3 Pyramidal Cell slow gamma PPC during VR periods | 15 - 79 | 18 - 101 |
| 2D, left | CA3 interneuron slow gamma PPC during flicker periods | 3 - 12 | 1 - 15 |
| 2D, right | CA3 interneuron slow gamma PPC during VR periods | 2 - 8 | 1 - 5 |
| 3D | Propsective coding ratio over position per day | 21 - 107 | 42 - 102 |
| 3E | Reward-related prospective coding ratio per day | 21 - 107 | 42 - 102 |
| 3F | Reward-related prospective coding ratio by trial speed | 21 - 107 | 42 - 102 |
| 3G | Place cell rate maps | 21 - 107 | 42 - 102 |
| 3H | Reward-related place cells | 1 - 52 | 5 - 18 |
| 4C | Prospective coding ratio during reward zone SWRs | 21 - 107 | 42 - 102 |
| 4D | Decoded position during reward zone SWRs | 21 - 107 | 42 - 102 |
| 4E | Proportion of prospective reward zone SWRs | 21 - 107 | 42 - 102 |
| 5D | Prospective coding ratio during fast and slow trials | 21 - 107 | 42 - 102 |
| 5E | Post-reward prospective coding | 21 - 107 | 42 - 102 |

**Supplementary Table 5.** Range of cells per animal included in main figure analyses.

| Supplementary Figure | Analysis | Cells per animal (min-max) | |
|---|---|---|---|
| | | Random | 40 Hz |
| 2B, CA3 PYR | Number of classified putative pyramidal cells | 24 - 68 | 31 - 100 |
| 2B, CA3 IN | Number of classified putative interneurons | 3 - 12 | 2 - 15 |
| 2B, CA1 PYR | Number of classified putative pyramidal cells | 13 - 79 | 18 - 101 |
| 2B, CA1 IN | Number of classified putative interneurons | 2 - 8 | 0 - 5 |
| 4C, left | CA3 pyramidal cell theta PPC during flicker periods | 8 - 67 | 28 - 99 |
| 4C, right | CA3 pyramidal cell theta PPC during VR periods | 23 - 66 | 22 - 73 |
| 4D, left | CA3 interneuron theta PPC during flicker periods | 3 - 12 | 1 - 15 |
| 4D, right | CA3 interneuron theta PPC during VR periods | 3 - 12 | 1 - 15 |
| 4E, left | CA3 pyramidal cell medium gamma PPC during flicker periods | 8 - 67 | 28 - 99 |
| 4E, right | CA3 pyramidal cell medium gamma PPC during VR periods | 23 - 66 | 22 - 73 |
| 4F, left | CA3 interneuron medium gamma PPC during flicker periods | 3 - 12 | 1 - 15 |
| 4F, right | CA3 interneuron medium gamma PPC during VR periods | 3 - 12 | 1 - 15 |
| 6A | Place cells | 21 - 107 | 42 - 102 |
| 6B | Place cells, spatial information | 21 - 107 | 42 - 102 |
| 6C | Place cells, sparsity | 21 - 107 | 42 - 102 |
| 6D | Place cells, peak position | 21 - 107 | 42 - 102 |
| 6E | Place cells, peak firing rate | 21 - 107 | 42 - 102 |
| 6F | Place cells, mean firing rate | 21 - 107 | 42 - 102 |
| 6G | Theta sequence strength | 21 - 107 | 42 - 102 |
| 6H | Prospective coding ratio over position per trial | 21 - 107 | 42 - 102 |
| 6I | Prospective coding ratio in the reward related zone | 21 - 107 | 42 - 102 |
| 6J | Propsective coding ratio in control vs. reward-related zone | 21 - 107 | 42 - 102 |
| 6K | Propsective coding ratio during high and low anticipatory licking trials | 21 - 107 | 42 - 102 |
| 6L | Decoding of current position | 21 - 107 | 42 - 102 |
| 6M | Decoding error of current position | 21 - 107 | 42 - 102 |
| 7B | Prospective coding ratio during all position SWRs | 21 - 107 | 42 - 102 |
| 7C | Proportion of prospective all position SWRs | 21 - 107 | 42 - 102 |
| 7G | Place cell activation | 21 - 107 | 42 - 102 |
| 7H | Place cell coactivation | 21 - 107 | 42 - 102 |
| 8B | Prospective coding ratio (both groups) on unengaged and engaged trials | 51 - 69 | 22 - 57 |
| 8C | Prospective coding ratio during unengaged and engaged trials | 51 - 69 | 22 - 57 |
| 8D | Prospective coding ratio difference between engaged and unengaged trials | 51 - 69 | 22 - 57 |
| 9A | Unoccupied zone decoding during theta | 21 - 107 | 42 - 102 |
| 9E | Unoccupied reward zone decoding by speed | 21 - 107 | 42 - 102 |
| 10A | SWRs with unoccupied reward zone decoding | 21 - 107 | 42 - 102 |

**Supplementary Table 6.** Range of cells per animal included in supplementary figure analyses.
